# Hydroxycinnamic acid amides emerge as multifunctional molecules involved in regeneration and volatile signalling during wound responses in tomato

**DOI:** 10.64898/2026.03.20.713112

**Authors:** C Grech-Hernández, C Andrade, Francisco Vera-Sirera, Ismael Rodrigo, José M Bellés, M Pilar López-Gresa, Purificación Lisón

## Abstract

- Hydroxycinnamic acid amides (HCAAs) are phenylpropanoid-derived metabolites with known antimicrobial and structural roles in plant defence against pathogens. However, their contribution to mechanical wound responses remains unclear, especially in terms of tissue regeneration and signalling.
- Here, we used tomato transgenic plants overexpressing the tyramine hydroxycinnamoyl transferase (THT), the key biosynthetic enzyme for HCAA production, to investigate the role of HCAAs in wound-induced responses, combining targeted metabolite profiling, gene expression, confocal microscopy, antioxidant assays, and volatile analyses.
- We show that *THT* overexpression enhances wound-induced accumulation of HCAAs, promoting vascular lignification, suberization, callose deposition, and increased regeneration capacity. Additionally, *35S::THT* plants display a distinct VOC profile that modulates defence gene expression in neighbouring wild-type plants, even in the absence of injury.
- These results identify *THT* as a key regulator of structural reinforcement and defence priming after mechanical damage. Our findings highlight a novel role for HCAAs in wound healing and interplant signalling, with potential applications for improving crop resilience to mechanical stress.

## Introduction

Plants, as sessile organisms, are constantly exposed to diverse environmental challenges, including mechanical injuries caused by herbivory, adverse climatic conditions such as wind, hail, and frost, or anthropogenic activities (Wasternack et al., 2006; Vega-Muñoz et al., 2020). These wounding events initiate a complex network of physiological, biochemical, and molecular responses that aim to mitigate damage, restore tissue integrity, and ensure survival. The effectiveness of these responses is not only critical for immediate recovery but also crucial for long-term adaptation and reproductive success.

The wound response begins with the perception of mechanical damage and the detection of damage-associated molecular patterns (DAMPs), which initiate downstream signalling cascades (Eulgem, 2000). These cascades are characterized by rapid increases in intracellular calcium levels, production of reactive oxygen species (ROS), and activation of transcriptional networks. Key phytohormones such as jasmonic acid (JA) coordinate local and systemic defence responses and tissue repair mechanisms (Wasternack et al., 2006). JA regulates the activation of defence genes and the production of secondary metabolites that deter herbivores and combat necrotrophic pathogens (Dammann et al., 1997). JA biosynthesis is initiated in the chloroplast with the oxygenation ofα-linolenic acid by lipoxygenases (LOX), followed by subsequent enzymatic steps involving allene oxide synthase and allene oxide cyclase, ultimately leading to JA production in the peroxisome (Li et al., 2018; Li et al., 2021). In addition, abscisic acid (ABA) has also been implicated in wound signalling. While traditionally associated with abiotic stress responses, ABA modulates wound-induced signalling and can influence JA biosynthesis (Wei et al., 2021). The interplay between JA and ABA is highly context-dependent and can be synergistic or antagonistic, thereby shaping the transcriptional output and amplitude of the defence response (Anderson et al., 2004). This hormonal crosstalk enables the plant to trade-off defence activation against growth and developmental priorities. In tomato, this balance is refined by the classic wound peptide systemin, which is perceived by the receptors SYR1 and SYR2 whose recently uncovered antagonistic functions modulate the extent and duration of the wound response (Zhou et al., 2025). Beyond hormonal signalling, wounding triggers the propagation of chemical, electrical and hydraulic signals from the damaged site to distal tissues, enabling systemic wound responses that prime the plant for future stresses (León et al., 2001). Among these signals, glutamate has been identified as a key wound signal that triggers long-distance calcium-based signalling via glutamate receptor-like channels, coordinating systemic defence responses across undamaged tissues (Toyota et al., 2018). This coordinated signalling cascade culminates in the production of late defence effectors, such as protease inhibitors (PINs), which accumulate both locally and systemically. These proteins interfere with the ability of herbivores to digest nutrients thereby reducing nutrient absorption and limiting further damage and represent a terminal step in the wound-induced defence program (Wasternack et al., 2006).

A key component of the plant defence arsenal is the production of secondary metabolites - terpenoids, alkaloids, and phenolics-which are pivotal in mediating responses to biotic and abiotic stresses (Reshi et al., 2023). Among them, terpenoids constitute the largest and most diverse group and play a central role in the composition of volatile organic compounds (VOCs) emitted by plants. These compounds serve as antimicrobial agents, insecticides, and herbicides (Pott et al., 2019), and many of their volatile members mediate ecological interactions by attracting beneficial insects such as pollinators and natural enemies of herbivores (Ninkuu et al., 2021). Beyond their direct protective roles, VOCs mediate indirect defence mechanisms by attracting the natural enemies of herbivores (Reshi et al., 2023) and facilitate plant-plant communication. In particular, VOCs emitted by stressed plants can prime defence responses in neighbouring plants, extending protective effects beyond the initially affected tissue or plant. Through this multifaceted repertory, plants sense, respond and adapt to also contribute to the plant defence system, function as herbivore deterrents and contribute to the therapeutic potential of plants (Bhambhani et al., 2021). Phenolics are involved not only in defence but also in plant development with the formation of structural polymers such lignin, suberin, and tannins, which enhance tissue strength and resistance to mechanical or environmental damage, including drought or wounding (Cesarino et al., 2022).

One class of phenolics with emerging significance in the defence response is the hydroxycinnamic acid amides (HCAAs). These are a group of low-molecular-weight phenylpropanoids formed by condensation of hydroxycinnamoyl-CoA thioesters with biogenic amines, such as aromatic β-phenylethylamines like tyramine or linear polyamines like putrescine. HCAAs play a dual role in plant defence. Structurally, they contribute to cell wall reinforcement, increasing resistance to both mechanical stress and pathogen attack (Clarke, 1982; Negrel and Lherminier, 1987). Functionally, HCAAs exhibit antimicrobial (Newman et al., 2001; Harris et al., 2010) and antioxidant activities, and some specific compounds like feruloyl-noradrenaline (*t*-FNA) show strong antioxidant properties and has been patented for its potential applications (López-Gresa et al., 2010). In Arabidopsis, HCAAs are known to accumulate in response to pathogen infection, contributing to callose deposition, especially coumaroyl-tyramine (CT) and coumaroyl-tryptamine (CTr), which are rapidly induced upon infection with *Pseudomonas syringae* and *Erwinia carotovora* (Macoy et al., 2022).

HCAAs are the products of an important branch of the phenylpropanoid pathway, in which phenylalanine ammonia lyase (PAL), cinnamic acid 4-hydroxylase, and 4-coumaric acid-CoA ligase are all involved (Liu et al., 2022). The final step in HCAA biosynthesis is catalysed by hydroxycinnamoyl-CoA:amine N-hydroxycinnamoyl transferase (THT) (Facchini et al., 2002). In tomato, THT is encoded by a multigene family, but only the isoform *tomTHT1-3*-hereafter referred to as *THT*-is strongly and rapidly induced during infection with avirulent *Pseudomonas syringae* pv. *tomato* (Pst), correlating with the selective accumulation of *p*-coumaroyl-noradrenaline and indicating a specialized role for this isoform in activating defence response (von Roepenack-Lahaye et al., 2003). Upon Pst infection tomato plants also exhibit increased *p*-coumaroyldopamine (CD) and feruloyldopamine (FD) (Zacarés et al., 2007) as well as other noradrenaline- and octopamine-derived amides such as *p*-coumaroylnoradrenaline (CNA), feruloylnoradrenaline (FNA), *p*-coumaroyloctopamine (CO), and feruloyloctopamine (FO) (López Gresa et al., 2011). Moreover, transgenic plants overexpressing *THT* (*35S::THT*) show constitutive accumulation of coumaroyl-tyramine (CT) and feruloyl-tyramine (FT), not only in leaves but also in flowers and fruits. Pathogen challenge in these plants leads to even higher levels CT and FT and amides derived from octopamine and noradrenaline, along with increased salicylic acid (SA) biosynthesis, stronger induction of pathogenesis related (*PR*) genes, and enhanced resistance to the bacterial pathogen (Campos et al., 2014). Studies have shown that HCAAs derived from tyramine contribute to a ligno-suberin vascular barrier that restricts *Ralstonia solanacearum* spread in resistant tomato varieties and enhance resistance by both mechanical reinforcement and antimicrobial activity (Kashyap et al., 2022). Although mechanical wounding has been shown to induce the accumulation of CT and FT (Pearce et al., 1998), recent reviews have established HCAAs as key components of plant responses to biotic and abiotic stresses (Xue et al., 2025), while underscoring their functional significance in wound-induced signalling and tissue repair remains a major unresolved question. In this context, the present study aims to investigate the response of tomato transgenic plants overexpressing THT to mechanical wound damage. By elucidating the function of HCAAs in mechanical damage defence, healing and tissue regeneration, this work contributes to a better understanding of how plants integrate metabolic, structural, and signalling strategies to defend against environmental challenges.

## Materials and Methods

### Plant material and growth conditions

In this study, we used tomato (*Solanum lycopersicum*) transgenic plants *35S:THT*, (*35S::THT)* overexpressing *THT* (Campos et al., 2014), and its corresponding parental line (WT), in Money Maker background. For the sterilization a mixture of sodium hypochlorite:distilled H_2_O (1:1) was used and sequential washings of 5, 10, and 15 min for the total removal of hypochlorite were performed. Germinated seeds were placed in 12-cm-diameter pots with vermiculite and peat. All procedures were performed under controlled greenhouse conditions with a 16/8 h light/dark photoperiod, approximately 50% relative humidity, and a temperature of 26 °C.

### Wounding and sample collection

After 4 weeks of growth, when the seedlings had developed 4-5 true leaves, five wounds were made along the veins in each of the five leaflets of the third and fourth leaves using toothed forceps, ensuring damage to both the abaxial and adaxial surfaces. Unwounded leaflets from the same leaves of control plants were collected as reference samples.

Leaf tissue samples were harvested 24 hours after wounding (24 hpw). The collected material was placed in 10 mL plastic tubes, immediately flash-frozen in liquid nitrogen, and stored at −80 °C until further analysis. The frozen tissue was subsequently homogenized under continuous cooling using a Retsch™ MM 400 mixer mill.

Simultaneously, leaf material for microscopy was also collected 24 hpw. Two wounded leaflets from the third and fourth leaves were excised and fixed in Falcon® tubes containing FAE buffer (50% absolute ethanol, 5% glacial acetic acid, 3.7% formaldehyde, and Milli-Q water) for subsequent processing and analysis.

### RNA isolation and RT-qPCR analysis

RNA extraction and cDNA synthesis from tomato leaves were performed using a silica membrane-based column kit (Macherey-Nagel GmbH, Germany) according to the manufacturer’s instructions. cDNA was synthesized from 1 µg of RNA using the PrimeScript RT reagent kit (Perfect Real Time, Takara Bio Inc., Otsu, Shiga, Japan) following the provided protocol. Quantitative PCR (qPCR) was conducted as previously described by Campos et al. (2014). Reactions were carried out in a 96-well plate with a final reaction volume of 10 µL per well. SYBR® Green PCR Master Mix (Applied Biosystems) was used as the fluorescent dye, and the actin gene served as the endogenous reference gene. All oligonucleotide sequences used in this study are listed in Table S1.

### Quantification of phenolic compounds by LC-MS

For the quantification of phenolic compounds, two types of extractions were performed: soluble and cell wall-bound phenolics. For both, 100 mg of frozen ground plant tissue were weighed into an Eppendorf tube. Regarding soluble ones, a modified version of López-Gresa *et al*. (2011) was used. The sample was extracted using 1 mL of 80% methanol (MeOH) containing 1 ppm of genistein as an internal standard. The mixture was incubated at room temperature for one hour to ensure efficient extraction. After incubation, the sample was centrifuged at 10,000 × *g* for 10 minutes. The supernatant was carefully transferred to a 4-mL vial. To maximize extraction efficiency, 1 mL of the same 80% MeOH solution containing genistein was added to the remaining pellet, followed by another centrifugation step under identical conditions. The supernatant obtained from this second extraction was pooled with the initial extract.

For the extraction of phenolic compounds bound to the cell wall, the protocol described by Santiago et al. (2018). The pellet from the previous centrifugation was dried for 24 hours at 37°C, and then 1.6 mL of 2N sodium hydroxide (NaOH) containing 1 ppm of genistein was added to each tube. The mixture was vortexed vigorously to break the pellet, followed by sonication in an ultrasonic bath for 20 minutes to enhance pellet disintegration. The tubes were then incubated at 65°C in a thermoblock for 10 minutes and vortexed again to ensure complete dissolution. Atmospheric oxygen was displaced by introducing nitrogen gas into the tubes, and the samples were incubated overnight at 37 °C in the dark with continuous agitation. The next day, the samples were centrifuged at 3,000 × *g* for 10 minutes. The supernatant was carefully transferred to a clean 4-mL glass vial, and the pH was adjusted to 2 by adding 600 µL of 6 N hydrochloric acid (HCl). Phenolic compounds were extracted by adding 1.6 mL of ethyl acetate to the acidified solution and vortexing the tube to promote phase separation. The upper organic phase was carefully collected and transferred to a clean 4 mL glass vial. This step was repeated twice, and the organic phases were pooled.

Once the extractions of the soluble phenolic compounds and the cell wall-bound phenolic compounds were completed, the vials corresponding to each extraction were dried under a stream of nitrogen gas. The residues were then resuspended in 100% methanol, filtered through a 0.22 µm membrane filter, and prepared for liquid chromatography analysis. Chromatographic separation and quantification were performed using an Orbitrap Exploris 120 mass spectrometer coupled with a Vanquish UHPLC System (Thermo Fisher Scientific, Waltham, MA, USA). LC was carried out by reverse-phase ultraperformance liquid chromatography using an Acquity PREMIER BEH C18 UPLC column (1.7 µm particle size, 2.1 × 150 mm; Waters Corp., Milford, MA, USA). The mobile phase consisted of 0.1% formic acid in water (phase A) and 0.1% formic acid in acetonitrile (phase B). The chromatographic gradient was programmed as follows: 0.5% solvent B over the first 2 min, 0.5-30% solvent B over 25 min, 30-100% solvent B over 13 min, 2 min at 100% B, a return to the initial 0.5% solvent B over 1 min, and re-equilibration at 0.5% B for 2 min. The flow rate was set at 0.4 mL/min, and the injection volume was 2 µL. The column temperature was maintained at 40°C. Ionization was performed using a heated electrospray ionization (H-ESI) source operating exclusively in negative mode. Data were acquired in full-scan mode at a resolution of 120,000 (FWHM). Compound identification was carried out using authentic standards synthesized according to (Zacarés et al., 2007). All chromatographic and mass spectrometric data were processed using TraceFinder software (Thermo Fisher Scientific, Waltham, MA, USA).

### Quantification of volatile organic compounds by GC-MS

For VOC analysis, 100 mg of homogenized tomato leaf tissue was placed in a 10 mL glass vial. To optimize VOC extraction, 1 mL of 6 M CaCl_2_ and 100 µL of 750 mM EDTA (pH 7.5) were added. The vials were airtight sealed and sonicated. VOCs were extracted using headspace solid-phase microextraction (HS-SPME) with a PDMS/DVB fibre as described by López-Gresa et al. (2017). The compounds were desorbed by heating the fibre. VOCs were analysed using an Agilent 8860 gas chromatograph (GC) coupled with an Agilent 5977B mass spectrometer (MS). Chromatography was performed on a 60 m, 0.25 mm ID silica column with a 5% phenyl/95% dimethylpolysiloxane stationary phase. Helium was used as the carrier gas.

GC-MS data were processed using MetAlign software, and the results were exported to Microsoft Excel for peak analysis and normalization. Quantification of VOCs was performed using Agilent MassHunter software, based on extracted ion chromatograms and peak integration. A One-Way ANOVA test was performed for each detected variable. Identification was based on mass spectra comparison with the NIST library and standard retention times. For multivariate statistical analysis, the normalized GC-MS dataset was further analysed using the MetaboAnalyst 5.0 web platform (www.metaboanalyst.ca). Ion peak intensities were autoscaled, and principal component analysis (PCA) was performed to explore global metabolic differences across genotypes and treatments. A heatmap was generated using the top 25 most discriminant ions, based on most positive loading plot ions of the first principal component and based on ANOVA (p < 0.05). Clustering was based on Euclidean distance and Ward’s linkage method, and visualization parameters included red/yellow/blue colour contrast and average group profiles. Six biological replicates per condition were included in the analysis.

### Hormone quantification

Quantification of plant hormones was performed using 100 mg of fresh leaf tissue per sample, and all results were normalized to fresh weight. Samples were homogenized and extracted with 80% methanol containing 1% acetic acid and a mixture of internal standards. After shaking for 1 hour at 4 °C, extracts were stored overnight at −20 °C. The supernatant was recovered by centrifugation, evaporated to dryness under vacuum, and resuspended in 1% acetic acid. For quantification of jasmonic acid (JA), abscisic acid (ABA), salicylic acid (SA), gibberellins (GAs), indole-3-acetic acid (IAA), and cytokinins (CKs), extracts were purified using Oasis HLB (reverse-phase) columns as described by Seo et al. (2011). For CKs, an additional purification step was performed using Oasis MCX (cation-exchange) columns, and the basic fraction containing cytokinins was eluted with 60% methanol and 5% ammonium hydroxide.

Separation of phytohormones was conducted using an Ultra-High-Performance Liquid Chromatography (UHPLC) system equipped with an Accucore RP-MS column (2.6 μm particle size, 100 mm × 2.1 mm; ThermoFisher Scientific). Hormones were separated using a linear gradient from 5% to 50% acetonitrile containing 0.05% acetic acid, at a flow rate of 400 μL/min over 21 minutes for JA, ABA, SA, IAA, and GAs. For cytokinins, a similar gradient was applied over 10 minutes. Hormone detection and quantification were performed using a Q-Exactive Orbitrap mass spectrometer (ThermoFisher Scientific) operating in targeted Selected Ion Monitoring (SIM) mode. Quantification was achieved using calibration curves embedded in the Xcalibur 4.0 and TraceFinder 4.1 SP1 software packages. Deuterium-labelled internal standards were used for each hormone, except for JA, for which dihydrojasmonic acid (dhJA) was employed.

### Antioxidant activity

For the antioxidant activity assay, 100 mg of plant material per sample was extracted using 500 μL of methanol (MeOH). following the procedure described by (Plazas et al., 2013) with minor modifications. After vortexing and centrifuging, the supernatant (Extract A) was recovered. The extract was then diluted with 100% ethanol to obtain a 2:3 dilution, referred to as Extract B. A 2,2-diphenyl-1-picrylhydrazyl (DPPH) solution was prepared by dissolving DPPH in absolute ethanol to make a 0.125 mM solution. For the reaction, 50 μL of extract B was mixed with 250 μL of 0.125 mM DPPH and 750 μL of absolute ethanol. The blank was prepared by replacing the extract with ethanol. The reaction was initiated by adding the DPPH solution to the samples and immediately measuring the absorbance at 517 nm using a TECAN microplate reader.

### Sample Fixation and Paraffin Embedding

Wounded areas were excised into 0.5 cm^2^ sections and immediately fixed overnight at 4 °C in FAE solution (50% absolute ethanol, 3.7% formaldehyde, 5% glacial acetic acid in Milli-Q water). Samples were subjected to two consecutive 5-minute vacuum infiltration cycles to enhance fixative penetration. After fixation, FAE was removed by sequential washes with ethanol solutions of gradually increasing concentrations. Subsequently, tissues were embedded in paraffin at 58 °C. Paraffin blocks were stored at 4 °C until sectioning. Transverse sections of the vascular bundles were obtained by cutting 10 μm thick sections using a microtome (Leica RM2255, Germany), and the sections were mounted onto standard microscope slides.

### Phloroglucinol Staining and Brightfield Microscopy

For lignin detection, paraffin sections were deparaffinized by two sequential incubations in Histoclear (7 and 5 minutes), followed by two washes in 100% ethanol (5 minutes each), two washes in 70% ethanol (5 minutes each) to remove eosin residues, and a final wash in 100% ethanol for 5 minutes. Phloroglucinol staining was performed immediately prior to imaging. A freshly prepared solution was used, consisting of 0.1 g phloroglucinol dissolved in 100 mL of 95% ethanol and mixed 1:1 with 37% hydrochloric acid. Sections were incubated in the staining solution for 5 minutes, briefly rinsed with 37% HCl, and mounted in 70% ethanol under a coverslip (Schindelin et al., 2013). Microscopic observation was conducted immediately after staining using a Leica DM5000 brightfield microscope. Images were acquired from transverse sections of vascular bundles, where lignified tissues appeared reddish to pinkish under transmitted light. Image acquisition and processing were performed using Fiji/ImageJ software.

### Basic Fuchsin and Auramine O Staining for Confocal Microscopy

For confocal imaging, wounded leaflets were clarified in ClearSee solution (10% xylitol, 15% sodium deoxycholate, and 25% urea in Milli-Q water) at room temperature under gentle agitation. The clarification process was maintained for approximately 2.5 weeks, with periodic replacement of the ClearSee solution whenever it became visibly green, to ensure optimal tissue clearing. Different leaf samples were used for each staining protocol, collecting tissues from the same region of the leaf.

For basic fuchsin staining, clarified samples were placed in 24-well plates containing 2.5 mL of staining solution per well (100 mg of basic fuchsin dissolved in 50 mL of ClearSee). After adding the staining solution, samples were subjected to two consecutive vacuum infiltration cycles of 5 minutes each to promote stain penetration. Samples were then incubated overnight at room temperature on a rocking platform. After staining, samples were washed three times in ClearSee for 20 minutes each under gentle agitation and were maintained in ClearSee until imaging (Ursache et al., 2018)

Auramine O staining was followed using a solution containing 250 mg of auramine O dissolved in 50 mL of ClearSee. Samples were vacuum infiltrated (two 5-minute cycles), incubated overnight at room temperature under gentle agitation, and washed twice in ClearSee for 20 minutes each. Following washing, samples were maintained in ClearSee until imaging (Ursache et al., 2018).

Imaging was performed by using a Leica TCS SP8 confocal microscope. For basic fuchsin, excitation was at 561 nm and emission was collected between 580-650 nm (pseudocolour red). For auramine O, excitation was at 488 nm and emission was collected between 505-550 nm (pseudocolour green). Sequential scanning was used to prevent signal overlap between channels. For each staining type, acquisition settings (laser power, gain, and offset) were kept constant across all samples to ensure comparability. Image acquisition was performed using LAS X software (Leica Microsystems), and images were subsequently processed with minimal adjustments to brightness and contrast.

### Callose staining and fluorescence quantification

To visualize callose deposition, plant tissue preserved in FAE buffer was subjected to vacuum infiltration for 10 minutes using a vacuum pump. Following fixation, the excess FAE was removed, and the samples were transferred to 70% ethanol and stored at 4 °C for 24 hours. To remove chlorophyll, tissues were sequentially treated with increasing concentrations of preheated ethanol (85%, 95%, and 100%) at 30-minute intervals. The final 100% ethanol step was repeated three times until complete decolorization was achieved. Ethanol was then replaced with sodium phosphate buffer, in which samples were incubated for 30 minutes at room temperature. The buffer was subsequently removed and replaced with a 0.05% aniline blue solution prepared in the same buffer. Samples were incubated overnight in darkness under gentle agitation (Currier, 1957). Finally, the staining solution was refreshed, and after 1 hour of additional incubation under the same conditions, samples were observed using a Leica 5000 UV microscope. Fluorescence intensity was quantified using ImageJ software.

### Tissue regeneration assays

Regeneration capacity was assessed in WT and *35S::THT* tomato seedlings using hypocotyl and cotyledon-based assays. Seeds were surface-sterilized and germinated in vitro on Murashige and Skoog (MS) medium, under controlled environmental conditions (16 h light/8 h dark photoperiod at 25°C). For hypocotyl regeneration, 7-day-old seedlings were transversely sectioned at the upper hypocotyl region using a sterile scalpel. Approximately 1-cm-long segments were laid horizontally on fresh MS medium, ensuring full contact between the explant and the medium surface to promote callus induction. For cotyledon regeneration, cotyledons from 7-day-old seedlings were excised and placed with the abaxial side facing the MS medium surface. Explants were incubated for 15 days under the same growth conditions. At the end of the incubation period, callus formation was manually scored for each explant under a stereomicroscope. A total of 20 explants per genotype were evaluated in each biological replicate, and the presence or absence of visible callus tissue was recorded.

### Tissue healing assays

To analyse leaf healing capacity, mechanical wounds were applied to fully expanded leaves of 4-week-old greenhouse-grown plants. Circular microperforations was generated in the lamina of the third and fourth true leaves using a 0.35 mm biopsy punch. Wounded leaves remained attached to the plant and were monitored *in situ*.

High-resolution images of each wound site were acquired at 7 and 14 days post-wounding (dpw). ImageJ software was used to quantify the ratio between wound area and perimeter, as an indicator of tissue contraction and healing progression. For this purpose, a macro command was developed to automate the analysis. Initially, the background was subtracted using a 2-pixel radius. A colour threshold was then applied (RGB: Red 0-255, Green 0-254, Blue 15-255) to isolate the wound area, and a binary mask (black and white) was generated. The binary “close” function was subsequently used to better define the healing ring. The perimeter of the ring and the internal area were measured to calculate the wound area-to-perimeter ratio. Comparative analyses were performed between WT and *35S::THT* genotypes using at least three independent biological replicates with a minimum of six wounds per condition.

### Inter-plant communication

For the inter-plant communication assays, methacrylate hermetic chambers (120 L) were used to establish controlled cohabitation environments, ensuring a sealed and uniform atmosphere. Five emitter plants (e-plants), consisting of either WT or *35S::THT* plants, subjected to wounded or non-wounded conditions, were placed within each chamber. Mechanical wounding of the emitter plants was performed as previously described, using toothed forceps to apply five wounds along the veins of five leaflets on the third and fourth leaves. Non-wounded emitter plants served as controls. Two healthy WT plants (r-plants) were used as receivers and cohabited in the same chamber alongside the emitter plants for a period of 24 hours. Following cohabitation, leaf tissues from the receiver plants were harvested, immediately flash-frozen in liquid nitrogen, and stored at −80 °C until further analysis (Pérez-Pérez et al., 2024).

As a positive control for volatile-mediated communication, three young WT plants (four weeks old, up to the fourth developed leaf) were exposed to 5 µM methyl jasmonate (MeJA) in additional chambers. MeJA was distributed onto several hydrophilic cotton balls placed within the chamber. They remained hermetically sealed for 24 hours and were opened only at the time of sample collection.

### Statistical Analysis

The analysis of variance was performed using the Two-Way ANOVA statistical test to determine whether there were significant differences between the different genotypes and conditions applied. Prior to this, the equality of variances was tested to assess whether the assumption of equal variances was valid for all the variables studied. If equal variances were assumed, ANOVA was conducted along with the corresponding Fisher least significant difference (LSD) post-hoc test. A *p*-value < 0.05 was considered statistically significant. Statistical treatments were conducted using Minitab 19 for Windows.

## Results

### *THT* overexpression selectively modulates wound-responsive gene expression without altering jasmonic acid and abscisic acid (ABA) levels

To investigate the role of *THT* in the tomato wound response, the *35S::THT* plants together with their parental line Money Maker wild type (WT), were characterized upon mechanical damage stress. Both genotypes were subjected to mechanical wounding, and samples were collected 24 hpw. Subsequently, the relative expression levels of several genes involved in wound signalling, including *THT, PAL5, LOX-D*, and *TCI21*, were quantified by qRT-PCR in both WT and *35S::THT* tomato plants under wounded and unwounded conditions (Fig. 1).

**Figure 1.**
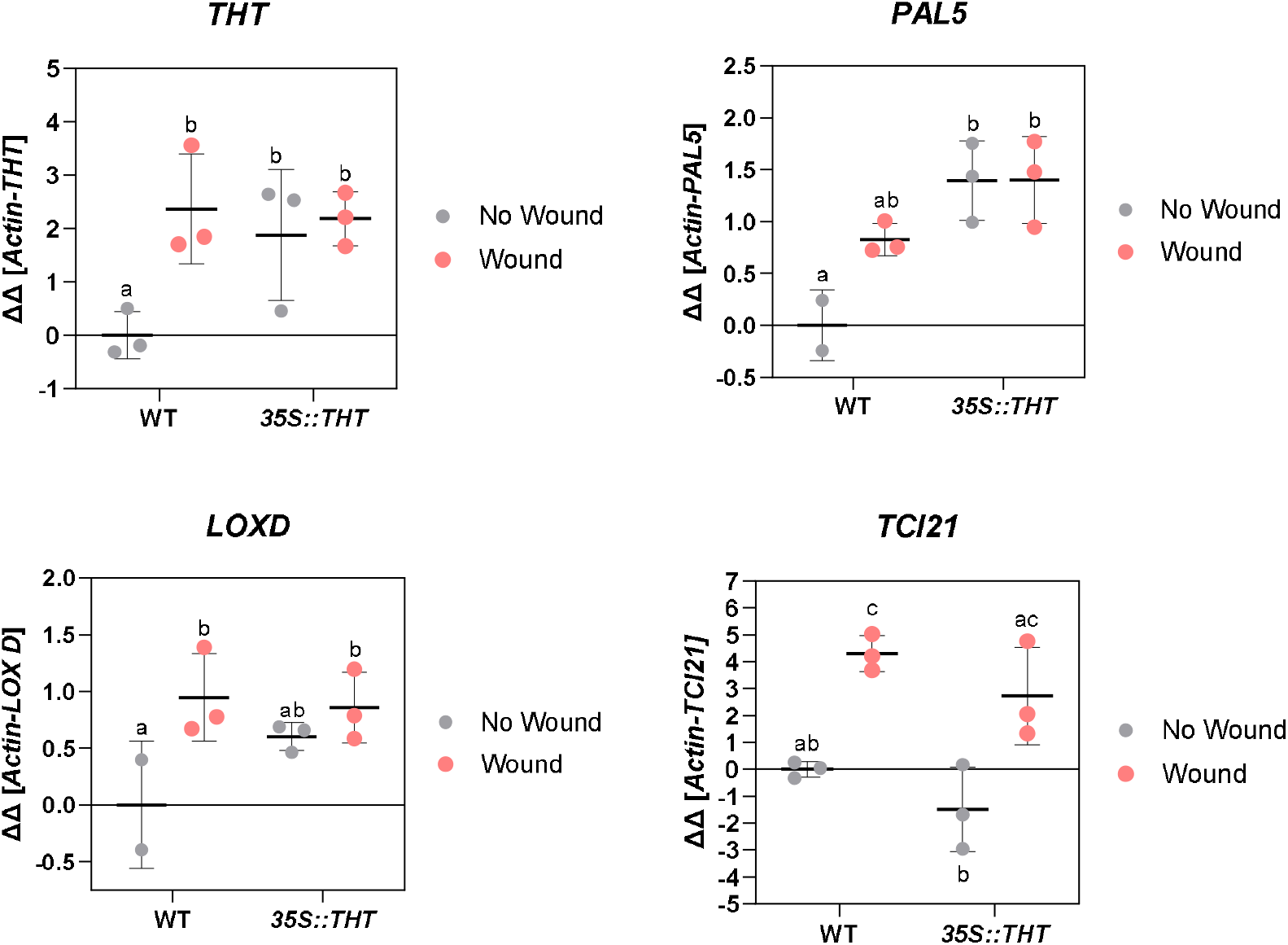
Relative expression of **A)** *THT*, **B)** *PAL5*, **C)** *LOX-D*, **D)** *TCI21* in leaves of transgenic tomato plants *35S::THT* and their parental Money Maker under non-wounded (No Wound) and wounded (Wound) conditions. Samples were taken after 24 hours of wounded plants (Wound) and without wounding (No Wound). The RT-qPCR values were normalized with the level of expression of Actin gene. The y axis represents the value of the Ct increment (ΔΔCt). The expression levels correspond to the mean ± SD of a representative experiment (n = 3). Statistically significant differences (ANOVA, *p* < 0.05) between genotypes and Wound or No Wound plants are represented by different letters.

In WT plants, wounding induced a significant upregulation of *THT* expression (Fig. 1A). In contrast, the *35S::THT* line exhibited markedly higher constitutive expression of *THT* regardless of wounding, as expected from the 35S promoter-driven overexpression. This elevated basal level in the *35S::THT* line indicates that the *THT* gene is strongly expressed in the absence of external stimuli, as previously reported by Campos et al. (2014). As a result, no further upregulation of *THT* expression was detected upon wounding in the *35S::THT* line. A similar significant expression pattern was observed for the *PAL5* and *LOX-D* (Fig. 1B, 1C), which are involved in the synthesis of phenylpropanoids (Guo and Wang, 2009) and jasmonic acid (JA; Yan et al., 2013), respectively. In WT plants, both genes showed a clear induction following wounding, whereas in the *35S::THT* plants, higher expression levels were observed even in the absence of wounding. This pattern indicates that while *PAL5* and *LOX-D* are wound-responsive in WT plants, their constitutively overexpression in the *35S::THT* line may reflect a pre-activated defence state. Notably, unwounded *35S::THT* plants exhibited significantly elevated *PAL5* expression compared to their parental WT line. Interestingly, the *35S::THT* plants did not show further induction of these genes upon wounding. Likewise, *LOX-D* is wound-inducible associated with the JA burst (Yan et al., 2013); however, its high basal expression in uninjured *35S::THT* tissues limits any further induction upon wounding.

Regarding *TCI21* (Fig. 1D), encoding a jasmonate-regulated proteinase inhibitor implicated in herbivore defence and a common marker for JA-mediated responses (Lisón et al., 2006), showed significant wound-induction in both *35S::THT* plants and their genetic background. However, no significant differences were detected in the basal (non-wounded) expression levels of this gene between the WT and *35S::THT* plants.

To check if the observed pre-activated level of expression in wound signalling genes could be due to changes in the hormonal content in the *35S::THT* line, we measured their JA and ABA levels, in the same samples (Fig. S1A, S1B). In both genotypes, wounding induced a significant increase in jasmonic acid (JA) and abscisic acid (ABA) levels, consistent with their central roles in wound-induced signalling pathways (Kimberlin et al., 2022; Peña-Cortés et al., 1989). Although JA and ABA levels increased significantly upon wounding in *35S::THT* plants, their accumulation did not differ from that observed in wounded WT plants.

These results indicate that the *35S::THT* plants retain a fully functional hormonal response to wounding, with significant increases in both phytohormones upon injury, comparable to those observed in WT plants. This suggests that THT overexpression does not suppress jasmonate or abscisic acid signalling but may instead act in parallel with these pathways. In line with this, *TCI21* expression remained inducible by wounding in both genotypes, indicating that specific components of the wound response remain responsive in *THT*-overexpressing plants.

Overall, these findings support the idea of a selective modulation of the wound response in *35S::THT* plants, where the constitutive and wound-induced accumulation of HCAAs may contribute not only to structural reinforcement but also to the fine-tuning of hormonal signalling pathways involved in stress adaptation.

### Mechanical wounding triggers enhanced accumulation of both soluble and cell wall-bound HCAAs in THT-overexpressing plants

Since *THT* expression is also induced by mechanical wounding, we next examined whether this transcriptional response translated into changes in HCAA accumulation following injury. To this end, we quantified the soluble levels of six HCAAs-derivatives: coumaroyl-tyramine (CT), coumaroyl-octopamine (CO), coumaroyl-dopamine (CD), feruloyl-tyramine (FT), feruloyl-octopamine (FO), and feruloyl-dopamine (FD), in leaves collected 24 hours post mechanical stress from both *35S::THT* and WT tomato plants, along with non-wounded controls (Fig. 2A-F).

**Figure 2.**
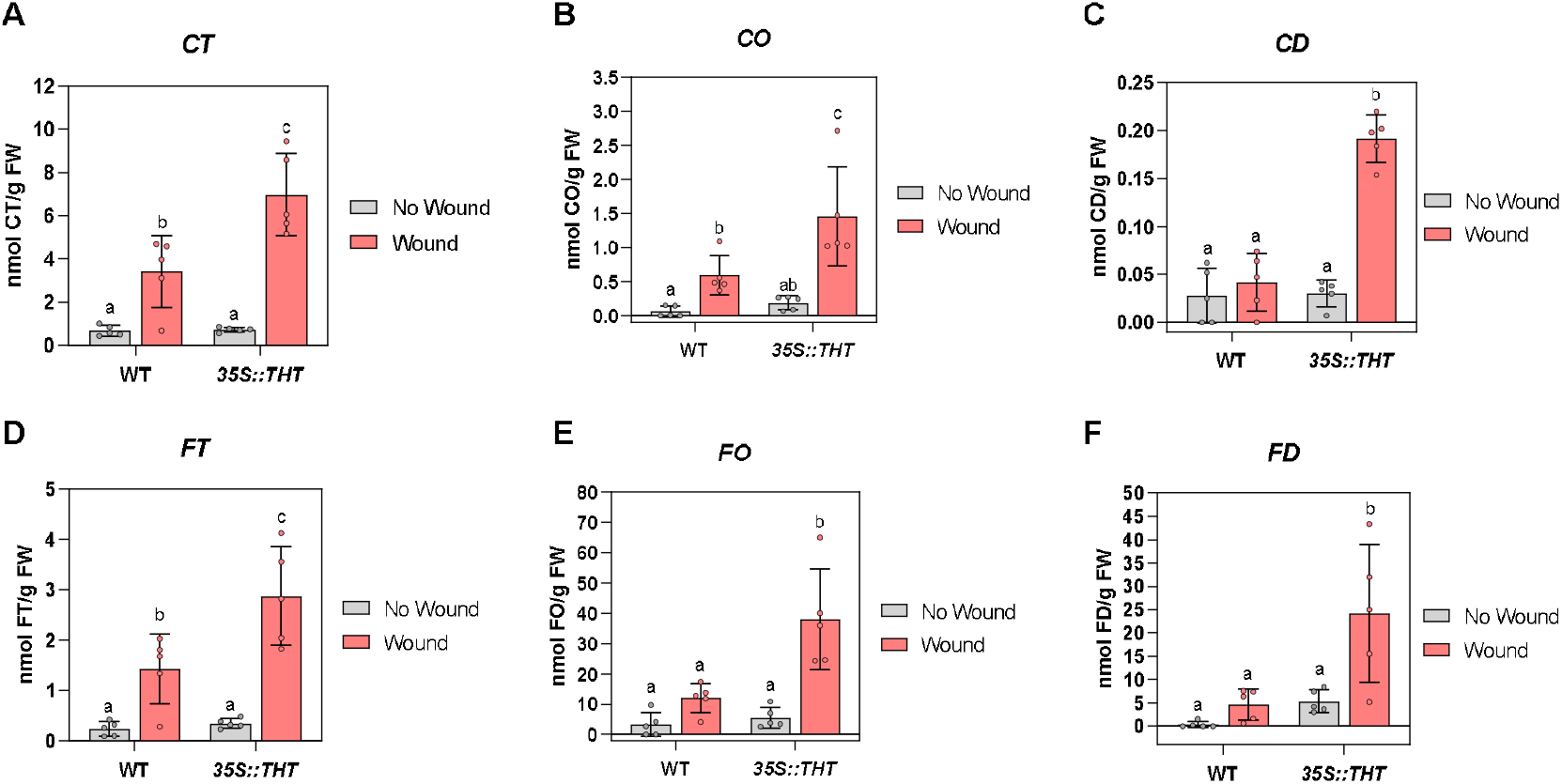
Levels of soluble hydroxycinnamic acid amides (HCAAs) in leaves of transgenic tomato plants (*35S::THT*) and their parental line Money Maker (WT) under non-wounded (No Wound) and wounded (Wound) conditions. Specifically, panels **A-C)** display coumaroyl derivatives: coumaroyl tyramine (CT), coumaroyl octopamine (CO), and coumaroyl dopamine (CD), and **D-F)** feruloyl derivatives, including feruloyl tyramine (FT), feruloyl octopamine (FO), and feruloyl dopamine (FD). Soluble HCAAs levels correspond to the mean ± SD of a representative experiment (n = 3). Statistically significant differences (ANOVA, *p* < 0.05) between genotypes and Wound or No Wound plants are represented by different letters.

In WT plants, wounding triggered a significant induction of CT, CO, and FT at 24 hpw (Fig. 2A, 2B, 2D). No significant differences were observed for CD, FO, and FD in these plants (Fig. 2C, 2E, 2F) at this timepoint. In contrast, *35S::THT* plants displayed a significantly higher accumulation of all six HCAAs analysed upon wounding. Collectively, these results indicate that *THT* overexpression enhances the wound-induced metabolic response, supporting the role for *THT* in boosting inducible defence-related phenylpropanoid metabolism.

To evaluate if these higher levels of soluble HCAA lead to their incorporation into the cell wall matrix, insoluble HCAAs derivatives were quantified in both WT and *35S::THT* plants under wounding, being detected CT, CO, FT and FO (Fig. 3A-D). In WT plants, wounding triggered a significant increase in cell wall-bound FT (Fig. 3C), indicating that FT incorporation is part of the wound-induced cell wall reinforcement response. In contrast, *35S::THT* plants exhibited a broader and more robust response, showing significant increases in both CT and FT levels upon wounding (Fig. 3A, 3C), highlighting an enhanced capacity to incorporate these compounds into the cell wall in these transgenic plants. Although wounding did not significantly increase CO levels in either genotype, wounded *35S::THT* plants accumulated substantially higher amounts than wounded WT, indicating that THT overexpression enhances CO accumulation under stress. Additionally, despite the lack of statistically significant differences between genotypes under non-wounded conditions, *35S::THT* plants consistently exhibited a trend toward higher levels of all four compounds, which may indicate a constitutive reinforcement of specific tissues associated with enhanced basal cell wall fortification.

**Figure 3.**
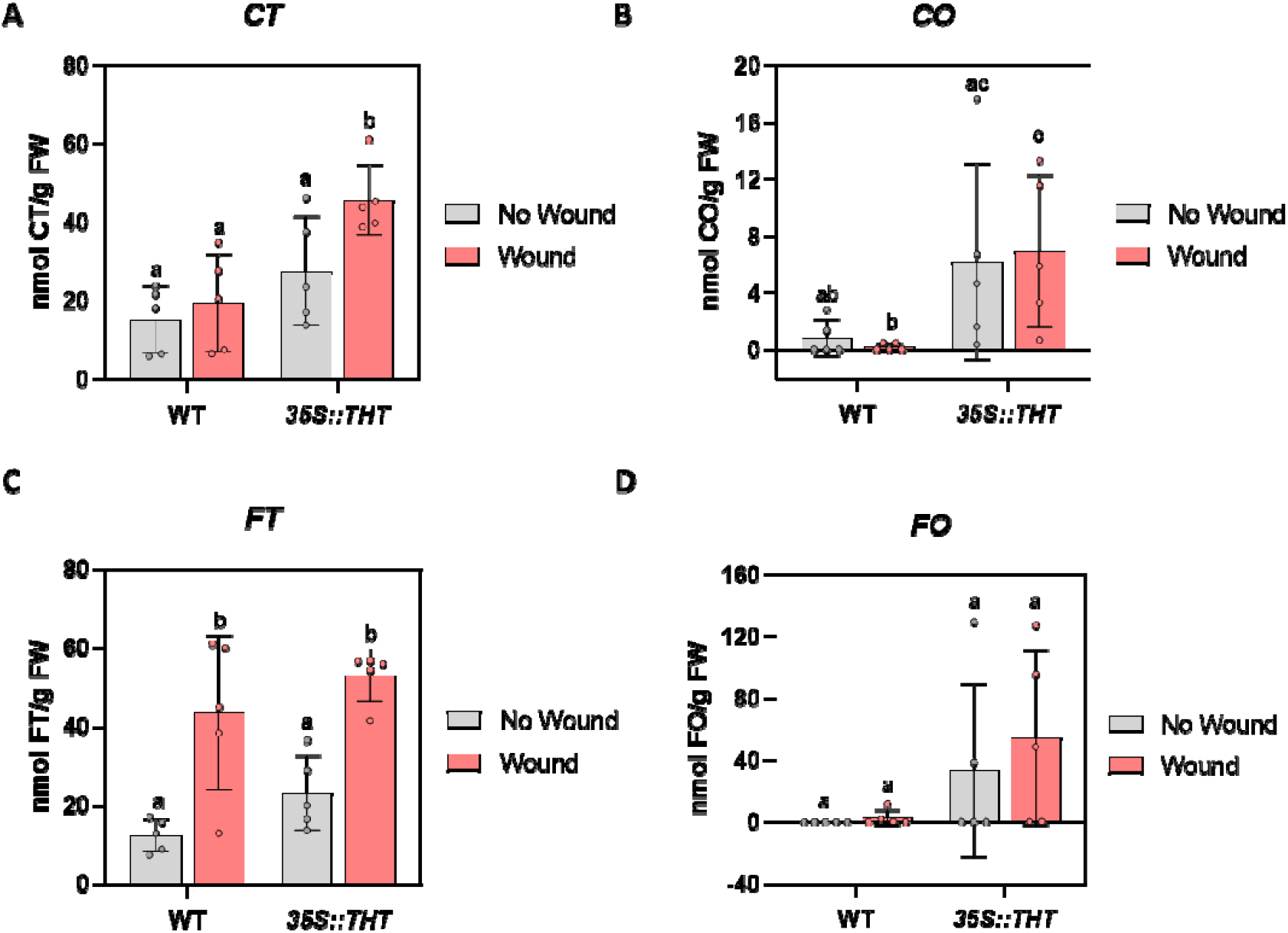
Levels of cell wall-bound hydroxycinnamic acid amides (HCAAs) in leaves of transgenic tomato plants (*35S::THT*) and their parental line, Money Maker (WT), under non-wounded (No Wound) and wounded (Wound) conditions. Panels A-C) show coumaroyl derivatives: coumaroyl tyramine (CT) and coumaroyl octopamine (CO), and C-D) feruloyl derivatives: feruloyl tyramine (FT) and feruloyl octopamine (FO). HCAA levels correspond to the mean ± SD of a representative experiment (n = 3). Statistically significant differences (ANOVA, *p* < 0.05) between genotypes and Wound or No Wound plants are represented by different letters.

Together, these results indicate that *THT* overexpression is associated with a general trend toward increased accumulation of cell wall-bound HCAAs, both under basal conditions and following wounding. This tendency suggests that *THT* may contribute to pre-existing structural differences between genotypesby enhancing the capacity to incorporate these compounds into the cell wall. Additionally, wounded WT plants also exhibited increased levels of specific HCAAs, supporting the role of cell wall reinforcement as part of the native wound response.

### THT overexpression enhances vascular lignification, suberin deposition and callose accumulation upon wounding

To explore whether the overexpression of *THT* affects structural reinforcement mechanisms in tomato leaves, we examined vascular tissue responses to mechanical wounding using histochemical and fluorescence-based staining. Since lignin, suberin, and callose are key components of secondary cell walls that contribute to mechanical strength and stress tolerance, we evaluated their spatial distribution and relative abundance in WT and *35S::THT* plants under control and wounded conditions. Our analyses focused on the vascular bundles and surrounding tissues to determine whether *THT* overexpression alters basal or wound-induced deposition patterns of these polymers.Mechanical wounding was applied to interveinal laminar regions of the leaf, deliberately avoiding direct damage to the main vein. To assess whether this localized injury triggered structural reinforcement in nearby vasculature, whole leaf preparations were stained with phloroglucinol-HCl to visualize lignin distribution. Strong lignin-associated signals were detected, localized exclusively within vascular bundles in the main vein of the leaf (Fig. 4A). In response to wounding, WT plants exhibited an increase in staining intensity and a noticeable thickening of the vascular bundles adjacent to the wound site. This response involved the reinforcement of pre-existing bundles through secondary wall thickening, without the formation of new vascular tissues. Notably, even in the absence of wounding, *35S::THT* plants exhibited thicker and more lignified vascular bundles compared with WT plants, suggesting that *THT* overexpression confers a basal state of structural fortification against mechanical stress. Examination of the regions directly subjected to mechanical damage revealed that the wounded area appeared more collapsed and showed greater tissue flattening in WT plants, indicative of increased susceptibility to applied pressure. In contrast, the wounded region displayed less compression and better preservation of structural integrity in *35S::THT* plants, despite being subjected to the same wounding method. This difference suggests that enhanced basal lignification in *35S::THT* plants contributes to greater local mechanical resistance to injury.

**Figure 4.**
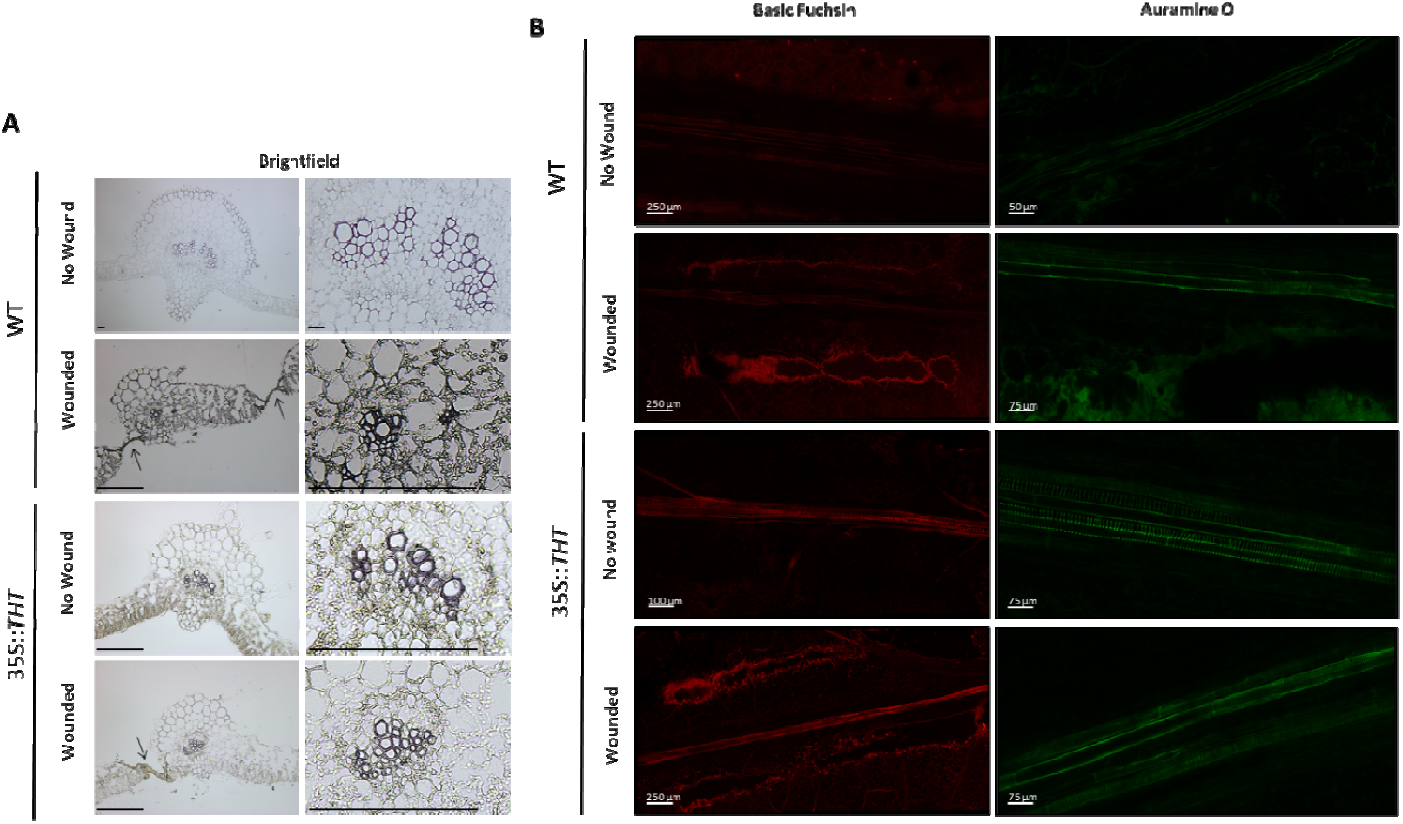
Histological and confocal analysis of cell wall modifications in wild-type (WT) and *35S::THT* tomato plants upon wounding. A) Brightfield microscopy images of leaf cross-sections stained with phloroglucinol–HCl to detect lignin. Transverse sections from WT and *35S::THT* plants under non-wounded (No wound) and wounded (Wound) conditions were collected 24 hours after wounding. Arrows indicate the wound site generated by pinching with forceps. Scale bars are indicated in each panel. B) Confocal laser scanning microscopy of leaves stained with basic fuchsin (red, lignin; excitation at 561 nm, emission 580–650 nm) and auramine O (green, suberin; excitation at 488 nm, emission 505– 550 nm). Images show vascular tissues from WT and *35S::THT* plants under No wound and Wound conditions. Scale bars are indicated in each panel. Images are representative of three independent biological replicates.

Confocal laser scanning microscopy of intact leaves stained with basic fuchsin and auramine O further confirmed these observations (Fig. 4B). In WT plants, minimal lignin fluorescence was detected under non-wounded conditions, whereas *35S::THT* plants displayed clear and continuous lignin signal along the vascular strand, indicating constitutive lignification. Upon wounding, lignin accumulation was observed around the wound site in both genotypes, but strong and continuous lignification along the vascular tissues was maintained only in *35S::THT* plants, highlighting their enhanced and spatially extended lignification response.

In parallel with the lignin pattern, suberin fluorescence was scarcely detected in non-wounded WT plants whereas it accumulated predominantly at the injury site upon wounding. In contrast, *35S::THT* plants exhibited suberin deposition in the vascular strand even in the absence of wounding, with further enhancement following mechanical damage.

Lignin and suberin signals were consistently overlapped within vascular tissues and wound-adjacent areas. This spatial convergence strongly suggests a coordinated deposition of both polymers as part of the reinforcement response. These results indicate that *THT* overexpression amplifies both basal and wound-induced structural reinforcement through coordinated lignin and suberin deposition.

To investigate whether the constitutive expression of *THT* also affects other structural defence compounds like callose, we examined its deposition in wounded leaflets using aniline blue staining and fluorescence quantification (Fig. 5A). Leaves from WT and *35S::THT* plants were wounded with forceps and collected after 24 hpw for histochemical analysis. Microscopic observation revealed that callose accumulated more abundantly at the wound sites of *35S::THT* plants compared to WT. Quantification of fluorescence intensity from nine independent images per genotype confirmed a significant increase in callose levels in *35S::THT* plants (Fig. 5B), suggesting that *THT* overexpression may potentiate cell wall reinforcement upon injury.

**Figure 5.**
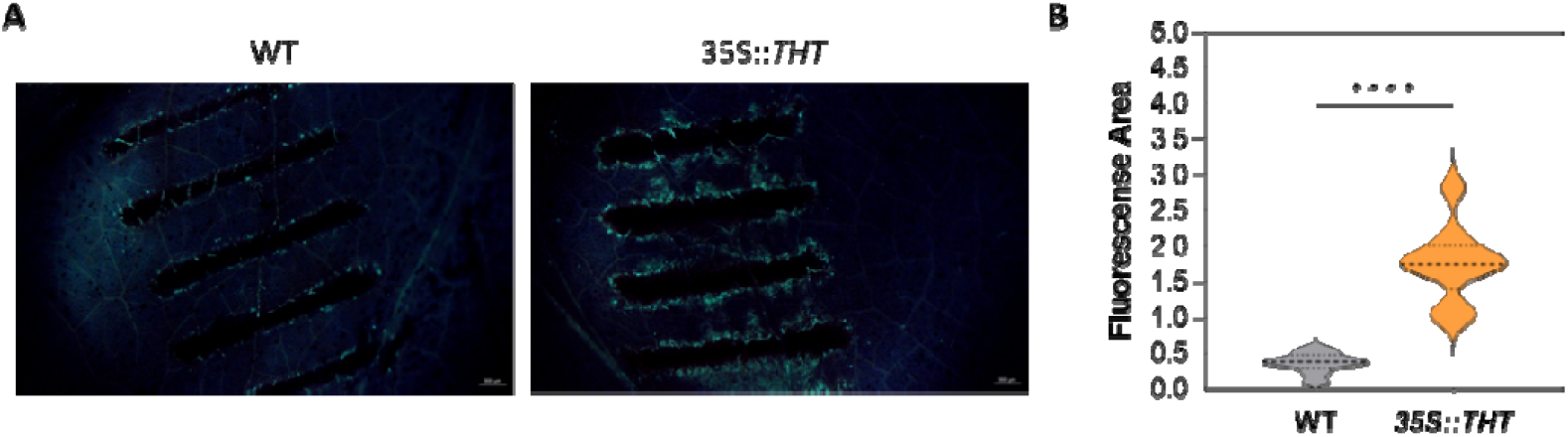
Callose deposition in wounded tomato leaves of WT and *35S::THT* plants. **A)** Aniline blue staining of wounded tomato leaves from *WT* and *35S::THT* plants. Callose deposition was detected 24 hours after wounding using a 0.05% aniline blue solution in sodium phosphate buffer. Samples were observed under a fluorescence stereomicroscope (*scale bar* = 300 μm). Three leaflets from independent plants per genotype were analysed, and three images were acquired per leaflet (*n* = 9). **B)** Violin-plot quantification of fluorescence intensity corresponding to callose deposition measured using ImageJ. Asterisks indicate statistically significant differences between genotypes (**** *p* < 0.0001; one-way ANOVA).

### THT overexpression confers a constitutively elevated antioxidant capacity in tomato plants

HCAAs are phenolic compounds with reported antioxidant capacity, which may contribute to the control of oxidative stress during wound responses. To explore the impact of mechanical damage on the antioxidant capacity of tomato plants, we monitored the DPPH radical scavenging activity in both WT and *35S::THT* plants under wounded and non-wounded conditions. Measurements were taken over a time course to monitor potential differences in the kinetics and magnitude of the antioxidant response between genotypes.

In WT plants, wounding led to a noticeable enhancement of antioxidant capacity compared to non-wounded controls (Fig. 6A). Troughout the time course, wounded WT plants consistently showed higher percentages of DPPH radical neutralization, with the difference becoming particularly evident after 30 minutes of reaction. This suggests that mechanical injury activates antioxidant defence mechanisms in WT plants, likely as a response to the oxidative burst associated with wounding. In contrast *35S::THT* plants exhibited a markedly different profile, showing a high basal antioxidant capacity under non-wounded conditions, comparable to that of wounded WT plants (Fig. 6A). Upon wounding, only a slight further increase in DPPH scavenging activity was observed, indicating that *35S::THT* plants possess an intrinsically enhanced antioxidant activity, highlighting the pre-activated defence status of the transgenic plants.

**Figure 6.**
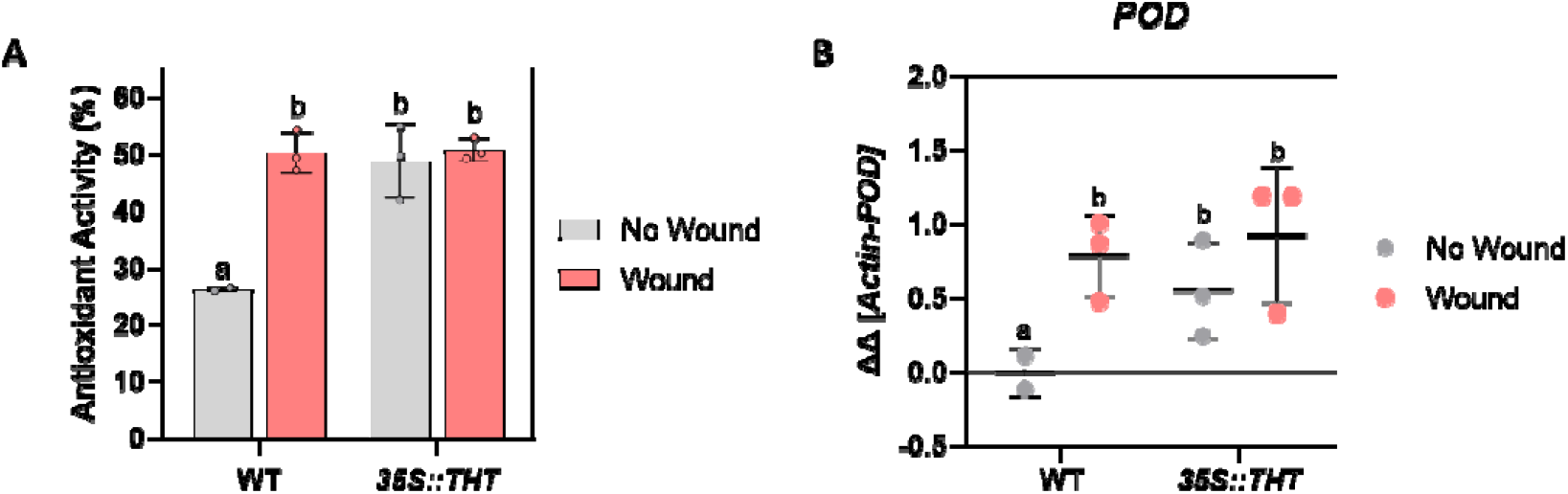
Antioxidant activity **A)** and expression of the oxidative stress-related gene *POD* **B)** in leaves of wild-type (WT) and transgenic tomato plants (*35S::THT*) under non-wounded (No Wound) and wounded (Wound) conditions. Samples were taken after 24 hours of wounding (Wound) and without wounding (No Wound). The RT-qPCR values were normalized with the level of expression of the Actin gene. The y axis represents the value of the Ct increment (ΔΔCt). The antioxidant activity and expression levels correspond to the mean ± SD of a representative experiment (n = 3). Statistically significant differences (ANOVA, *p* < 0.05) between genotypes and Wound or No Wound plants are represented by different letters.

To further investigate whether enzymatic antioxidant responses were associated with these observations, the expression levels of a peroxidase (*POD*) gene were analysed by RT-qPCR (Fig. 6B). The results revealed that *POD* expression was significantly induced in WT plants following wounding, whereas *35S::THT* plants maintained elevated levels of *POD* expression irrespective of wounding. This expression pattern closely mirrored the antioxidant capacity profiles observed in the DPPH assay, supporting the notion that *THT* overexpression contributes to a constitutive activation of antioxidant defences.

Overall, these findings suggest that overexpression of *THT* enhances tomato plant resistance to oxidative stress by maintaining a constitutively elevated antioxidant capacity therefore minimizing the need for further induction of antioxidant responses upon wounding.

### *THT* overexpression promotes wound healing and callus regeneration

To determine whether *THT* overexpression influences tissue recovery following mechanical injury, we assessed wound healing and callus regeneration in WT and *35S::THT* tomato plants using both *in planta* and *in vitro* approaches. Based on the previously described phenotypes like HCAAs accumulation, cell wall remodelling and structural and antioxidant differences between genotypes, we hypothesized that *35S::THT* plants may exhibit enhanced regenerative capacity. To test this, wound healing dynamics, callus induction efficiency, and endogenous hormone profiles were analysed to elucidate potential mechanisms underlying the observed phenotypes.

The effect of *THT* overexpression on wound healing was evaluated using a recovery leaf assay. Circular microperforations (0.35 mm diameter) were inflicted in leaves of WT and *35S::THT* tomato plants using a biopsy punch, and the ratio of healed tissue ring area and the original hole perimeter was measured at 7 and 14 days post-wounding (dpw). This ratio was used as an indicator of wound-edge expansion and tissue remodelling, as evidenced by its progressive increase in WT plants between 7 and 14 dpw, with higher values reflecting more advanced healing responses (Fig 7A). Notably, by 7 dpw, *35S::THT* plants already reached values comparable to those observed in WT at 14 dpw (Fig. 7A), indicating an accelerated wound healing response associated with *THT* overexpression.

**Figure 7.**
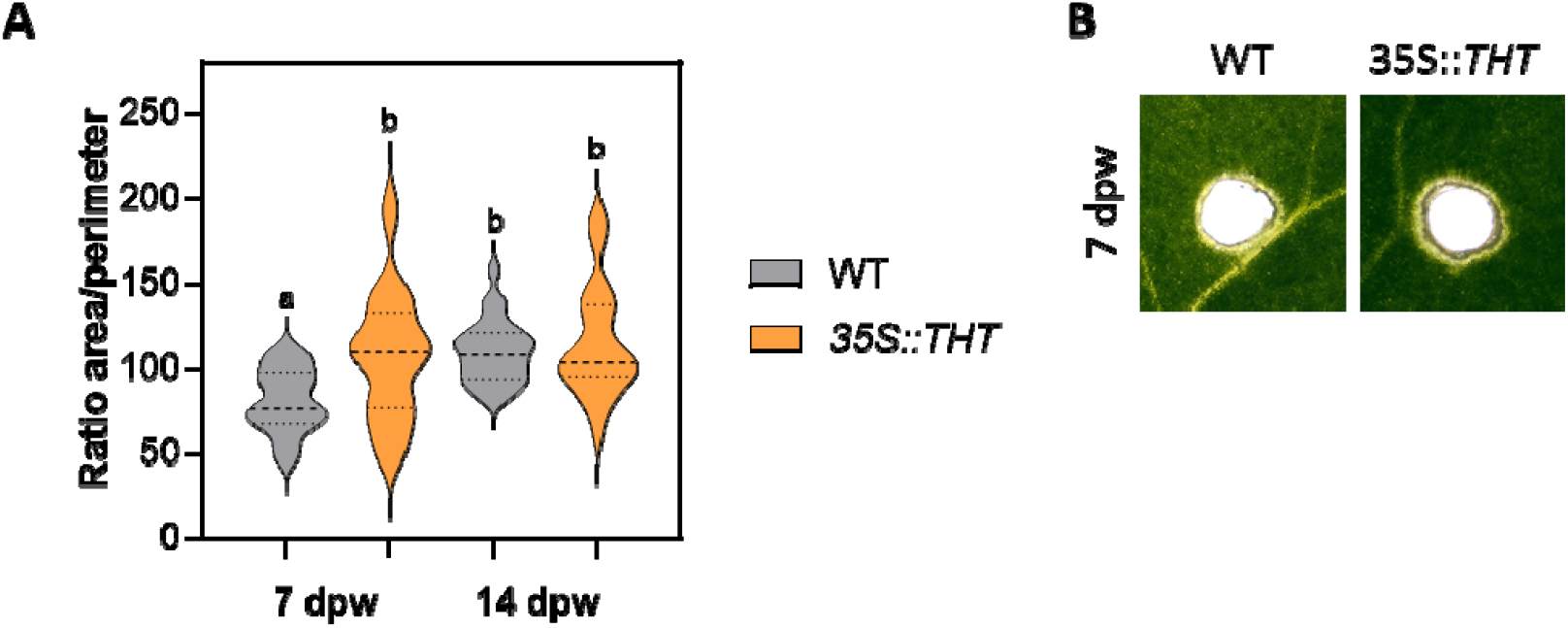
Wound healing dynamics in WT and *35S::THT* tomato plants. **A)** Ratio of area to perimeter in circular wounds made on the third and fourth leaves of 4-week-old greenhouse-grown WT and *35S::THT* tomato plants at 7-and 14-days post-wounding (dpw). **B)** Representative images of WT and *35S::THT* hypocotyls are shown, after 7 dpw. Statistically significant differences (ANOVA, *p* < 0.05) between genotypes and Wound or No Wound plants are represented by different letters.

To assess *in vitro* regenerative capacity, hypocotyl and cotyledon explants from 7-day-old seedlings were cultured on MS medium. In hypocotyl segments, the frequency of callus formation was significantly higher in *35S::THT* than in WT plants (Fig. 8A, 8C). A similar pattern was observed in cotyledon explants, where the proportion of callus-forming explants was also increased in *35S::THT* plants (Fig. 8B, 8D).

**Figure 8.**
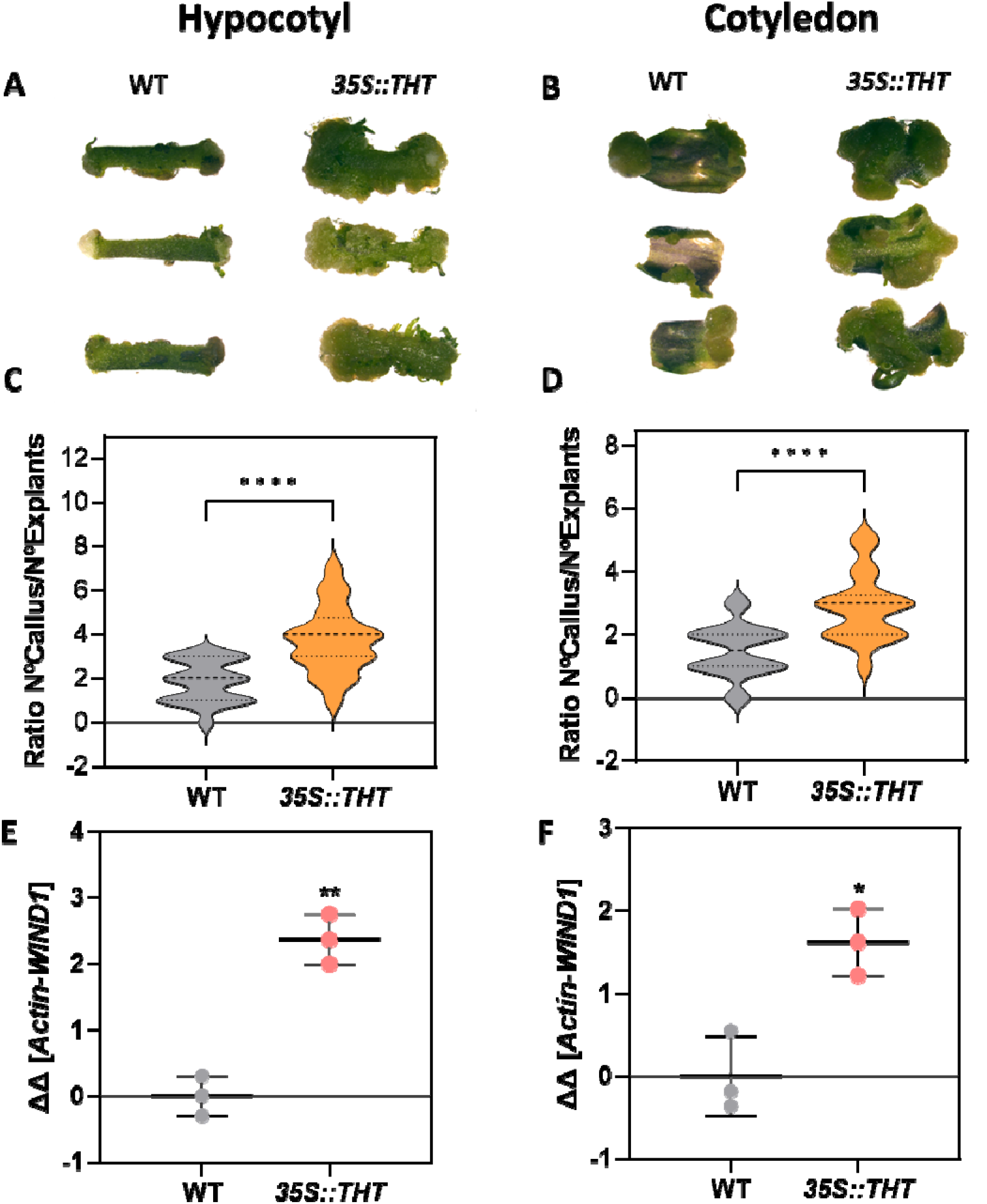
*In vitro regenerative capacity* in WT and *35S::THT* tomato plants. **A–B)** Representative images of callus formation from explants of 7-day-old seedlings after 15 days of in vitro culture. **A)** Hypocotyl explants and **B)** cotyledon explants from WT and *35S::THT* plants are shown. **C-D)** Callus formation ratio (Nº callus/Nº explants) in **C)** hypocotyl segments and **D)** cotyledon explants of 7-day-old seedlings cultured for 15 days on MS medium. E–F) Relative expression of the regeneration-associated gene *WIND1* in E) hypocotyl and F) cotyledon explants from WT and *35S::THT* plants. Explants were collected after 15 days of in vitro culture. RT–qPCR values were normalized to the expression level of the *Actin* gene. The y-axis represents Ct increment values (ΔΔCt) and correspond to the mean ± SD of a representative experiment (n = 3). Statistically significant differences (ANOVA, *p* < 0.05) between genotypes are indicated by asterisks. Data represent the mean ± SD of a representative experiment (n = 3).

To investigate the molecular basis of enhanced regeneration in our transgenic lines, we quantified the expression of *WIND1* (WOUND⍰INDUCED DEDIFFERENTIATION 1), a key AP2/ERF transcription factor known to mediate wound⍰induced cellular reprogramming and callus formation (Yang et al., 2024). Consistent with its proposed role in linking wound signals to regenerative pathways, *WIND1* expression was significantly upregulated in hypocotyl (Fig. 8E) and cotyledon (Fig. 8F) tissues of *35S::THT* plants when compared with WT controls, indicating a strong induction of this regeneration⍰associated gene in the overexpression background. This elevated *WIND1* expression in *35S::THT* correlates with the enhanced regenerative phenotypes observed in these tissues, supporting the hypothesis that *THT* overexpression promotes activation of transcriptional networks involved in cellular dedifferentiation and regeneration.

To investigate potential hormonal mechanisms underlying these phenotypes, endogenous levels of indole-3-acetic acid (IAA), dihydrozeatin (DHZ), and iso-pentenyladenine (iP) were quantified in leaves from wounded and non-wounded plants 24 hpw. Under basal conditions, *35S::THT* plants showed significantly higher IAA levels compared to WT, which remained unchanged upon wounding (Fig. S1C). In contrast, DHZ (Fig. S1D) and iP (Fig. S1E) levels were stable across genotypes and treatments, with no significant differences detected.

Taken together, these results indicate that *THT* overexpression promotes wound healing and tissue regeneration in tomato, potentially through modulation of auxin homeostasis and enhanced regenerative capacity.

### VOCs emitted by *THT* overexpressing plants differentially modulate defence gene expression in WT receiver plants

To investigate how volatile cues from genetically distinct plants influence defence activation in neighbours, we conducted interplant communication assays using closed methacrylate chambers. Wild-type (WT) receiver plants were exposed for 24 hours to VOCs emitted by emitter plants of either WT or *35S::THT* genotype, under non-wounded or wounded conditions. As a positive control, additional WT plants were treated with methyl jasmonate (MeJA).

Expression analysis of the jasmonate-responsive gene *TCI21* revealed no significant changes in WT receiver plants exposed to VOCs from wounded WT or *35S::THT* emitter plants, compared to exposure to non-wounded WT plants (Fig. 9A). However, exposure to non-wounded *35S::THT* emitters resulted in a significant increase in *TCI21* expression, suggesting that these transgenic plants emit defence-activating VOCs even in the absence of tissue damage. In contrast, the expression of *PR1*, a salicylic acid marker gene, showed a different pattern (Fig. 9B). WT receivers exposed to VOCs from WT wounded plants displayed a significant upregulation of *PR1* compared to those exposed to non-wounded WT plants. Interestingly, this expression level was comparable to that observed in plants exposed to non-wounded *35S::THT* emitter plants, indicating that *35S::THT* plants may constitutively release signals that mimic the effect of wounding in WT. However, exposure to wounded *35S::THT* emitter plants led to a decrease in *PR1* expression relative to the non-wounded *35S::THT* condition, suggesting that tissue damage in *THT*-overexpressing plants modifies the emitted VOC profile in a way that reduces salicylic acid-associated responses.

**Figure 9.**
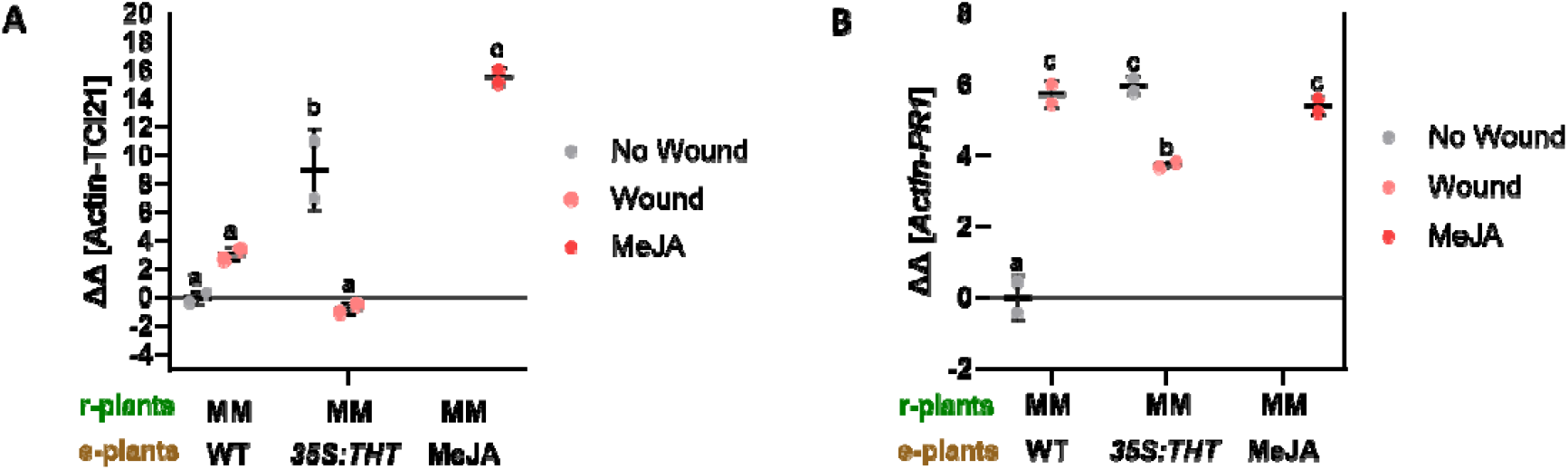
Analysis of interplant communication mediated by volatile compounds emitted by tomato plants undergoing wounding. Evaluation of the defensive response in wild-type Money Maker plants used as receivers (r-plants) after 48 hours of cohabitation with emitter plants (e-plants) of different genotypes and wounding conditions. Emitter plants included wild-type (WT) and transgenic *35S::THT* lines, both non-wounded (No Wound) and mechanically wounded (Wound), as well as plants exposed to a volatile methyl jasmonate treatment (MeJA, 50 μM) used as a positive control. Relative expression of **A)** *TCI21 and* **B)** *PR1* in r-plants after cohabitation. The RT-qPCR values were normalized with the level of expression of Actin gene. The y axis represents the value of the Ct increment (ΔΔCt). The expression levels correspond to the mean ± SD of a representative experiment (n = 3). Statistically significant differences (ANOVA, *p* < 0.05) between genotypes and Wound or No Wound plants are represented by different letters.

These findings indicate that *35S::THT* plants constitutively emit VOCs capable of modulating gene expression in neighbouring WT plants, but that their wounded state may produce a qualitatively different volatile signal that partially suppresses specific branches of the defence network.

### THT overexpression reshapes basal and wound-induced VOC emission profiles in tomato

To assess the influence of *THT* on the emission of volatile organic compounds (VOCs) in response to wounding, the volatilome of *35S::THT* tomato lines was compared to that of their wild-type (WT) parental line. Mechanical wounding was applied using toothed tweezers, and samples were collected 24 hours post-injury, as detailed in the Materials and Methods section. An untargeted metabolomic approach employing gas chromatography-mass spectrometry (GC-MS) was applied to identify and quantify VOC emissions following wound-induced stress.

To explore VOCs differences emissions between WT and *35S::THT* non wounded and wounded leaves a principal component analysis (PCA) was performed after 24 hpw (Fig. 10). The first principal component (PC1), accounting for 45.1% of the variance, distinguished the samples based on the wound-induced volatilome: wounded samples were positioned on the positive axis of PC1, while unwounded samples were on the negative axis. The second principal component (PC2), explaining 17.8% of the variance, segregated the samples by genotype: WT parental line plants were located on the negative axis of PC2, whereas the *35S::THT* plants were located on the positive axis of PC2. The higher PC1 percentage of variance supports that mechanical wounding effect exerted a major impact on VOC emissions, surpassing the effect of genotype.

**Figure 10.**
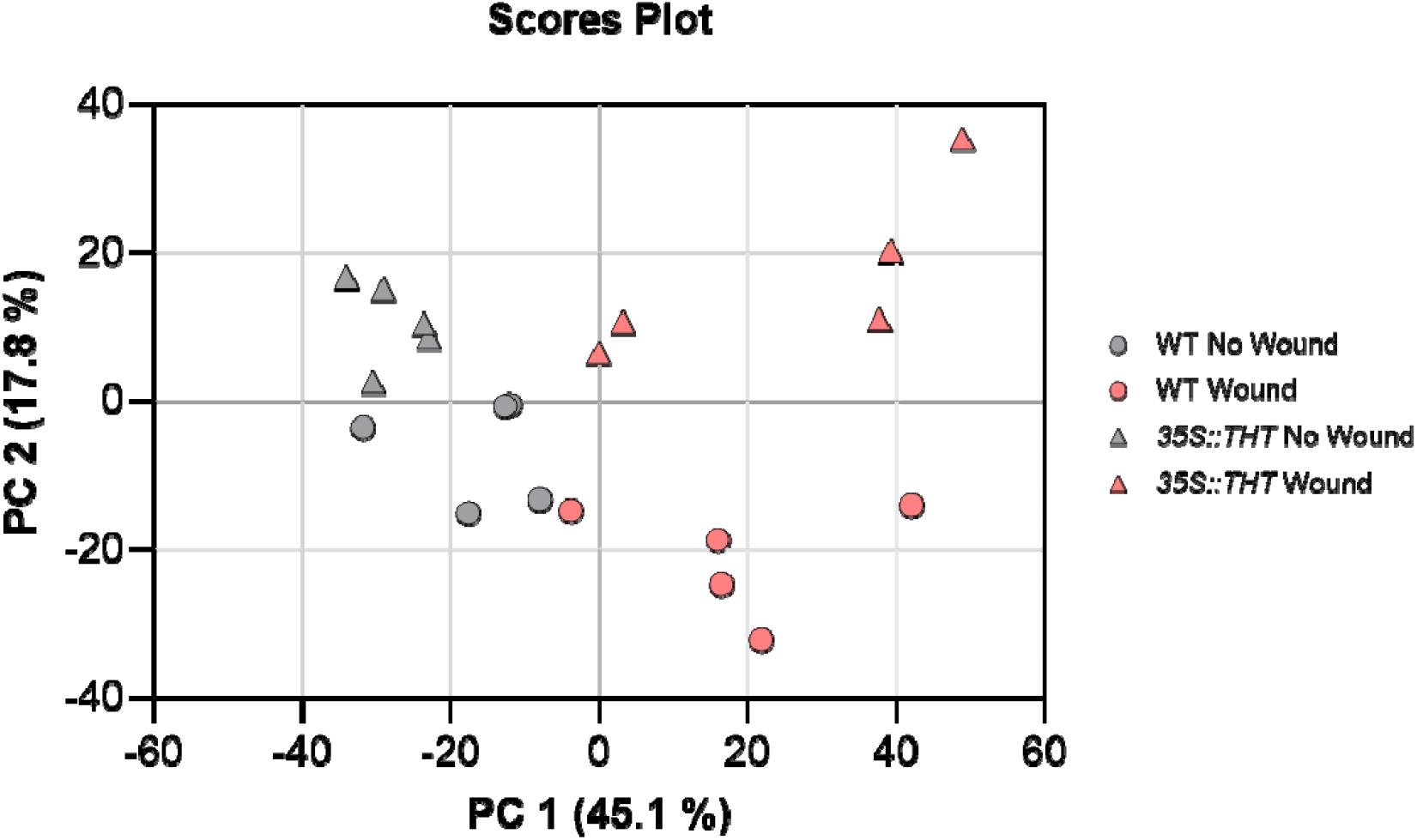
Score plot of PCA based on mass spectrum signal abundance within the m/z range of 35 to 250, corresponding to tomato leaf samples from WT and *35S:THT* plants under non-wounded (No Wound) and wounded (Wound) conditions. The data points are represented as circles for WT plants and triangles for *35S:THT* transgenic plants. The colours indicate the conditions: grey for non-wounded plants and pink for wounded plants. Principal component 1 (PC1) explains 45.1% of the total variance and Principal component 2 (PC2) accounts for 17.8% of the variance.

To identify the volatile compounds differentially emitted by *35S::THT* and WT plants after wounding, we performed a loading plot from the PCA analysis. Subsequently, a list of VOCs that most strongly contributed to the separation across PC1 and PC2 was extracted. We focused in the compounds with a higher positive value on PC1 and PC2 to identify the most exclusive ions from wounded *35S::THT* tomato plants, which included *E*-2-heptenal, *E*-2-octenal, *E*-2-decenal, *2*-pentylfuran, and 2H-pyran-2,6-dione, among others. This suggests that the overexpression of *35S::THT* alters the volatile plant response to wounding. Wounded WT plants emitted volatiles characterized by a higher positive loading value on PC1 and lower values on PC2, predominantly comprising classical mechanical stress volatiles, such as key green leaf volatiles (GLVs) and C5 volatiles, including 1-penten-3-one, *Z*-3-hexenal, *Z*-3-hexenyl butyrate (HB), and 2,4-hexadienal. Interestingly, plants overexpressing *THT* also displayed a distinct VOC profile under non-wounded conditions clustering in the negative region of PC1 and the positive region of PC2 in the PCA score plot. Among the volatiles differentially over-emitted by no wounded *35S::THT* leaves, analysis of the loading plot revealed several monoterpenes, including the monoterpenes sabinene, α-terpinene, *D*-limonene, and β-cymene, as well as the unsaturated aldehyde *E*-2-nonenal. These compounds were absent or present at much lower levels in the corresponding WT samples.

To further visualize the emission dynamics of these key volatiles across genotypes and wounding, a heatmap was generated using their relative abundance under all conditions (Fig. 11). This representation reveals distinct treatment and genotype specific VOC signatures, with three clearly defined clusters: (i) an aroma of wounding characterized by VOCs predominantly emitted by WT plants upon injury mostly defined by GLVs, (ii) an aroma associated with *THT* overexpression, marked by monoterpenoids, and iii) a wound-induced *THT* aroma, reflecting the synergistic activation of both wounding and *THT*-mediated metabolic reprogramming, emphasizing the unique volatile profile of the transgenic line under stress.

**Figure 11.**
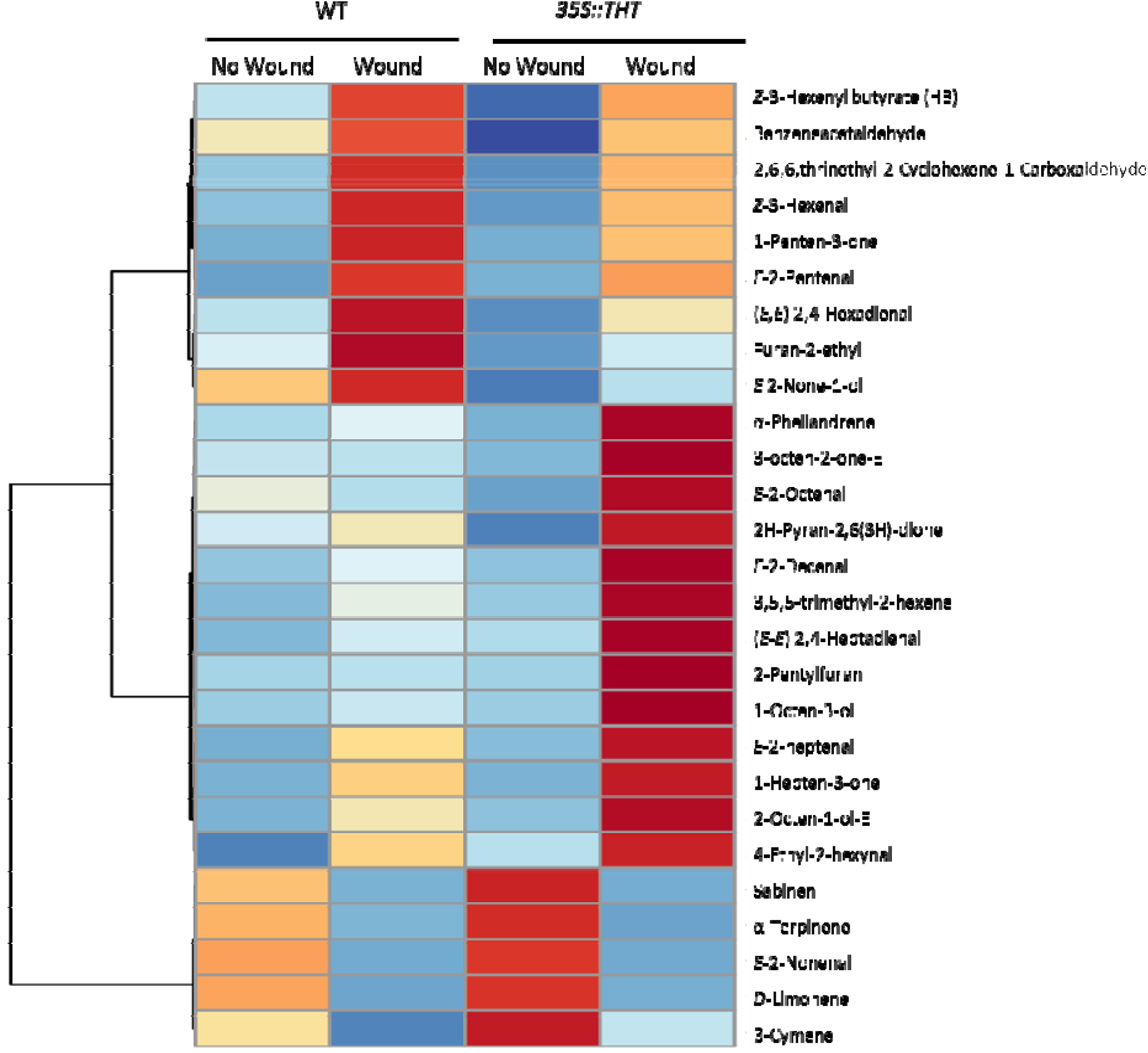
Distinct volatile profiles in *35S::THT* and WT plants. Heatmap of volatile compounds in *WT* and *35S::THT* tomato leaves under *No Wound* and Wound conditions. Volatile profiles were obtained from leaf samples collected 24 hours after mechanical wounding. Data represent the mean of six biological replicates per condition (*n* = 6). Compounds were identified based on retention time and mass spectral data, and are indicated by name. Clustering was performed using Ward’s linkage method and Euclidean distance. Colour scale indicates auto scaled values of relative abundance (red: high; blue: low).

### Volatile cues emitted by WT and THT-overexpressing plants differ in their defence signalling capacity

Based on the comparative volatile emission of WT and *35S::THT* plants, three representative compounds were selected for quantification and functional assays: *Z*-3-hexenal, predominantly emitted by wild-type plants (Fig. 12A), *E*-2-heptenal, enriched in wounded *35S::THT* plants (Fig. 12B) and limonene which displayed higher emission by both non-wounded transgenic lines (Fig. 12C).

**Figure 12.**
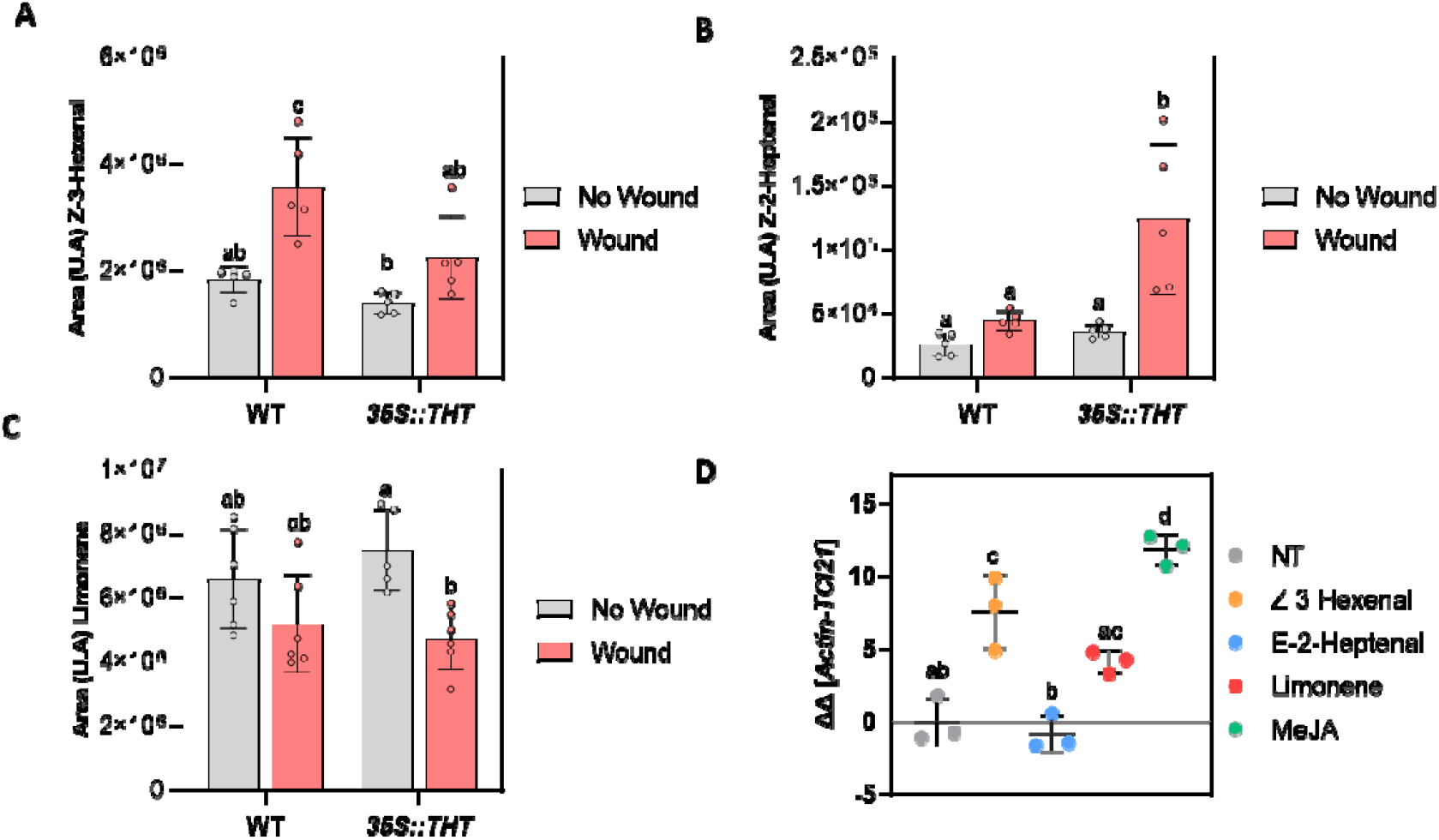
Quantification of selected volatile compounds and their effect on the expression of the jasmonate-responsive gene *TCI21*. **A-C)** Relative quantification of Z-3-hexenal, E-2-heptenal and Limonene in leaf samples of WT and *35S::THT* plants under wound and No Wound conditions. Data correspond to the mean ± SD of normalized signal areas from five biological replicates (n = 5). D) Relative expression of the jasmonate-responsive gene *TCI21* in WT plants treated exogenously with Z-3-hexenal, E-2-heptenal and limonene. A non-treated plant was used as negative control, and a plant treated with methyl jasmonate (MeJA) was included as positive control for *TCI21* induction. The RT-qPCR values were normalized with the level of expression of Actin gene. The y axis represents the value of the Ct increment (ΔΔCt). The expression levels correspond to the mean ± SD of a representative experiment (n = 3). Statistically significant differences (ANOVA, *p* < 0.05) between the treatments are represented by different letters.

To assess their potential as defence-activating signals, MM receiver plants were exogenously treated with these compounds, alongside MeJA as positive control, and the expression of *TCI21*, a jasmonate-responsive marker associated with wound response, was analysed (Fig. 12).

Among the volatiles tested, only *Z*-3-hexenal treatments significantly induced *TCI21* expression compared to untreated controls, supporting its role as an effective interplant signalling molecule capable of activating jasmonate-related defence pathways (Fig.12D) In contrast, exposure to *E*-2-heptenal did not alter *TCI21* expression levels (Fig. 12D), indicating that although this compound is enriched in wounded *35S::THT* plants, its perception does not require further activation of jasmonate-responsive defences, potentially reflecting the constitutively elevated defensive status of these plants. Furthermore, limonene exposure exhibited a tendency to increase *TCI21* expression, despite not reaching statistical significance (Fig. 12D). Notably, limonene is constitutively emitted by *35S::THT* plants in the absence of wounding, reinforcing the notion that these transgenic lines maintain a basal defence status that resembles the induced state of wounded wild-type plants. The lack of robust *TCI21* induction by treatment with limonene alone suggests that defence activation in receiver plants exposed to VOCs from non-wounded *35S::THT* plants is not mediated by a single compound, but rather by the combined action of a specific volatile blend. These observations suggest that the constitutive VOC aroma emitted by non-wounded *35S::THT* plants is selectively oriented towards the activation of jasmonate-related defence responses, as evidenced by the induction of TCI21 (Fig 9A), with limonene appearing as a potential mediator of this activation (Fig 12D). This constitutive volatile profile may contribute to a pre-activated signalling environment, further supporting enhanced readiness against potential threats.

## Discussion

Mechanical damage triggers tightly coordinated defence and regeneration responses in plants, but the metabolic nodes that connect structural reinforcement, redox control and tissue regeneration are still largely unresolved. In this work, we identify tyramine hydroxycinnamoyl transferase-derived hydroxycinnamic acid amides as a central metabolic hub integrating these processes during the tomato wound response.

### THT overexpression selectively rewires wound-associated transcriptional networks without altering core hormone signalling

Constitutive overexpression of the *THT* gene in tomato leads to a selective reprogramming of wound-associated gene expression (Fig.1) while preserving intact jasmonate and abscisic acid signalling (Fig.S1A, S1B). As expected, THT transcripts are strongly elevated under basal conditions in *35S::THT* plants, whereas its induction upon wounding occurred only in WT plants (Fig.1A). This pattern is consistent with previous studies in other plant species, where *THT* was shown to be a wound-inducible gene. In potato tuber discs, *THT* activity increased dramatically in the outermost cell layers after injury, remaining elevated for several days and contributing to the accumulation and integration of FT and FO into the cell wall matrix during tissue healing (Negrel et al., 1993). Similarly, in maize leaves, *THT* activity was clearly induced 3 hours after wounding, reaching a maximum at 36 hours, with levels 40 times higher than in unwounded tissues. This induction coincided with the accumulation of hydroxycinnamoyltyramines, particularly FT (Ishihara et al., 1999). These observations support a conserved role for *THT* in early wound responses across species.

Upstream of THT, *PAL5* expression (Fig. 1B) was significantly upregulated in *35S::THT* plants under basal conditions and remained unchanged after wounding. Induction of *PAL* in response to wounding has been documented in multiple plant species, including non⍰vascular (e.g. *Marchantia polymorpha*) and vascular plants such as lettuce, where PAL transcript accumulation occurs in tissues adjacent to the wound site (Campos et al., 2004; Yoshikawa et al., 2018). In tomato, our data identify *PAL5* as a wound-responsive isoform in WT plants, supporting its involvement in phenylpropanoid activation following mechanical damage.

The constitutive up⍰regulation of *PAL5* in *35S::THT* lines suggests a metabolic re⍰wiring that positions phenylpropanoid metabolism in a pre-activated state. Such a configuration may confer a selective advantage by reducing the latency between damage perception and metabolite biosynthesis, thereby ensuring rapid availability of hydroxycinnamic precursors for downstream HCAA formation. Given that tomato possesses a multigene PAL family, comprising several isoforms with potentially distinct regulatory and functional roles (Zhang et al., 2023), future work will be required to determine whether additional PAL isoforms contribute to this basal activation and whether specific isoforms are preferentially channelled toward HCAA production rather than other phenylpropanoid branches.

In contrast, for the JA-related genes *LOX-D* (Fig.1C) and *TCI21* (Fig. 1D), no significant differences were detected between genotypes, showing a clear induction in response to wounding and confirming their role as reliable markers of mechanical damage. The induction of *LOX-D* upon wounding aligns with its established function in the biosynthesis of JA, where it acts as a chloroplast-localized lipoxygenase rapidly upregulated by mechanical injury and systemic signals such as systemin and methyl jasmonate (Fritig et al., 1998). Consistently, functional studies of the tomato *spr8* mutant, which carries a point mutation in the catalytic domain of TomLoxD, confirm that TomLoxD is required for wound-induced JA biosynthesis and effective defence responses (Yan et al., 2013). The fact that both, *LOX-D* and *TCI21*, are responsive but not differentially expressed in *35S::THT* plants highlights the robustness of the JA signalling cascade, which remains intact despite alterations in phenolic metabolism.

This genetic behaviour, with no changes in *LOX-D* and *TCI21*, was mirrored at the hormonal level since no significant differences were observed in JA or ABA levels between *35S::THT* and wild-type plants, either under basal conditions or after injury (Fig.S1). This represents the first report examining JA and ABA levels in *35S::THT* plants, offering novel insight into how constitutive *THT* expression may affect the hormonal responses to mechanical stress. As reported previously, JA accumulates rapidly following wounding, often within the first hours, and is later followed by an increase in ABA levels. This temporal pattern suggests a coordinated response in which JA initiates early defence signalling, while ABA contributes to sustaining wound healing and limiting water loss (Lulai et al., 2009). In various plant systems, including Arabidopsis, poplar, and potato, JA and ABA have been shown to act synergistically to protect tissues under stress conditions and promote regeneration. JA enhances the early wound response and tissue reinforcement, while ABA supports long-term protection and recovery processes such as suberization and regeneration under stress (Wan et al., 2025; Xu et al., 2024). Together, our data indicate that THT overexpression does not interfere with canonical wound hormone pathways but instead acts in parallel to them.

### THT-mediated accumulation of hydroxycinnamic acid amides reinforces cell wall architecture

A central outcome of THT overexpression is the enhanced accumulation of HCAAs, such as CT and FT, both in soluble forms (Fig. 2A, 2D) and bound to the cell wall (Fig. 3A, 3C). Early work in tobacco established that THT activity is a key determinant of the rate and localization of wound-induced HCAA accumulation, promoting the rapid deposition of both soluble and cell wall-bound amides around damaged tissues (Hagel & Facchini, 2005). This phenotype extends previous observations in tomato plants, where *THT* overexpression resulted in elevated HCAA levels and enhanced resistance to pathogens such as *Pseudomonas syringae* (Campos et al., 2014), and is consistent with reports showing that ligno-suberin vascular coatings enriched in tyramine-derived HCAAs restrict *Ralstonia solanacearum* colonization in resistant tomato varieties (Kashyap et al., 2022).

HCAAs occupy a strategic position at the interface between metabolism and structure. When incorporated into the cell wall, they contribute to ligno-suberin assemblies that strengthen vascular tissues and restrict damage propagation and in their soluble form, they exhibit antimicrobial and antioxidant activities. This phenotype extends previous observations where *THT* overexpression resulted in elevated HCAA levels and enhanced resistance to pathogens such as *Pseudomonas syringae* (Campos et al., 2014). In addition, recent work in resistant tomato varieties has shown that the induced formation of vascular coatings enriched in ligno-suberin and HCAAs derived from tyramine restricts *Ralstonia solanacearum* colonization, acting as a physical and chemical barrier to limit pathogen spread (Kashyap et al., 2022). HCAAs are known to play crucial roles in plant defence mechanisms, acting as antimicrobial agents and contributing to cell wall reinforcement (Liu et al., 2022; Berti et al., 2025). The increased accumulation of these compounds in *THT*-overexpressing plants suggests that *THT* activity is a key factor in modulating the phenylpropanoid pathway, initiated by *PAL* induction (Fig.1B), towards the biosynthesis of defence-related metabolites.

Microscopic analyses support this dual structural role, revealing localized reinforcement at wound sites in *35S::THT* plants, particularly within vascular tissues, where increased lignin- and suberin-associated signals were detected (Fig.4). The elevated basal expression of *PAL5* in *35S::THT* plants indicates increased activation of the early steps of the phenylpropanoid pathway, which not only feeds HCAA biosynthesis but also provides precursors for lignin and suberin (Kashyap et al., 2022). Recent studies have shown that under stress conditions, such as vascular infections or mechanical injury, plants can activate combined lignification and suberization programs in response to hormonal and redox signals, in which phenolic amides may play a functional role (Lee et al., 2019; Somssich, 2020; Romero et al., 2021). Therefore, the combined evidence of HCAA accumulation, elevated *PAL5* expression, and structural rearrengements observed by microscopy in *35S::THT* plants supports the hypothesis that *THT* overexpression contributes to the structural preparation of tissues to contain or repair damage through the formation of reinforced physical barriers.

In addition to ligno-suberin reinforcement, transgenic *35S::THT* tomato plants exhibited a significantly higher accumulation of callose compared to their parental WT line, suggesting that *THT* overexpression enhances structural defence responses to injury (Fig. 5). This increased callose deposition around the wounded areas coincided with a greater accumulation of HCAAs in the transgenic plants, indicating a potential functional link between both processes. HCAAs such as CT and FT accumulate in damaged tissues and contribute to cell wall remodelling (Bassard et al., 2010; Macoy et al., 2022), potentially facilitating callose deposition as a compensatory and protective mechanism following wounding. Several studies have shown that exogenous application of HCAAs, such as CT and coumaroyltryptamine, can induce callose accumulation in Arabidopsis under biotic stress (Macoy et al., 2022). In this context, our results suggest that *THT* overexpression may increase the availability of HCAAs, thereby indirectly promoting callose biosynthesis as part of a generalized structural reinforcement mechanism in response to physical damage.

Thus, THT emerges as a key regulator of mechanical defence architecture in tomato, promoting both constitutive and wound-induced reinforcement of the cell wall through coordinated deposition of lignin, suberin and callose.

### Enhanced antioxidant capacity in *35S::THT* plants supports wound resilience

Analysis of antioxidant activity revealed that *35S::THT* plants exhibit significantly higher basal antioxidant capacity compared to WT plants, suggesting a constitutive activation of redox-regulatory mechanisms. Following wounding, both genotypes showed an increase in antioxidant activity, but *35S::THT* plants consistently maintained superior levels (Fig. 6), suggesting an enhanced ability to buffer oxidative stress associated with tissue damage.

This phenotype is consistent the accumulation of HCAAs, phenolic compounds with well-established antioxidant properties (López Gresa et al., 2011). Overexpression of *THT* likely enhances the biosynthesis of highly reducing HCAAs, such as FT and CT which could contribute to oxidative homeostasis and help mitigate cellular damage following wounding. Supporting this interpretation, HCAAs have been shown to accumulate in infected tissues during plant immune responses, where they reinforce cell walls and modulate ROS levels (Liu et al., 2022). Similarly, Macoy et al. (2022) showed that CT and coumaroyl tryptamine are induced in *Arabidopsis* upon pathogen challenge, acting as regulators of redox signalling. More broadly, Khawula et al. (2023) highlighted the dual role of hydroxycinnamic acid derivatives as modulators of oxidative stress responses and promoters of plant growth under adverse environmental conditions. In support of this redox-activated state, the expression of a peroxidase-encoding gene (*POD*) was also found to be constitutively upregulated in *35S::THT* plants, further suggesting an enhanced capacity for oxidative stress management.

Finally, acummulative evidence suggests that HCAAs operate at the interface between redox control and hormonal signalling. In tomato, THT overexpression has previously been linked to elevated salicylic acid (SA) levels following pathogen infection (Campos et al., 2014), supporting the idea that enhanced HCAA pools contribute to an activated defence state. Consistent with a link between SA signalling and HCAAs metabolism, tomato lines with constitutively high SA (SlS5H-silenced) show increased accumulation of HCAAs (Payá et al., 2022). Although the molecular mechanisms remain to be elucidated, such redox-hormonal crosstalk may represent an additional layer through which THT modulates wound resilience by integrating antioxidant protection with broader defence signalling pathways (Yang et al., 2019).

Recent work has shown that efficient wound regeneration relies on sustained redox homeostasis rather than on the suppression of oxidative stress. Ruiz-Solaní et al. (2025) demonstrated that bacterial cellulose-induced tissue regeneration depends on the coordinated activation of ROS production and antioxidant buffering, allowing redox-dependent regenerative processes to proceed without excessive cellular damage. Similarly, the constitutively elevated antioxidant capacity observed in *35S::THT* plants may favour wound resilience by maintaining ROS within a productive signalling range that supports tissue recovery following damage.

### Rebuilding after injury: role of HCCAs in tomato healing and regeneration

Our results show that *THT* overexpression significantly enhances leaf wound healing and tissue regeneration (Fig.8) in tomato. This effect was evidenced by accelerated wound closure in leaves *in vivo* (Fig.7) and increased callus formation from hypocotyl and cotyledon explants cultured *in vitro* (Fig. 8), hallmarks of active regenerative programs involving cell proliferation, tissue remodelling, and local genetic reprogramming.

The enhanced regenerative performance of *35S::THT* plants correlated with elevated basal IAA levels (Fig.S1C), suggesting that *THT* overexpression may modulate auxin homeostasis, possibly through interactions with the metabolism of HCAAs, which have been implicated in the regulation of hormone activity and cell division in other plant systems. Previous studies have shown that certain HCAAs, such as feruloyl amides, can influence auxin activity and the organization of regenerative tissue (Campos et al., 2014; Liu et al., 2022). In this context, we propose that the accumulation of HCAAs in *35S::THT* plants could alter local IAA dynamics, promoting a more permissive environment for regeneration. In contrast, no changes were detected in cytokinin levels (Fig. S1D, S1E), suggesting that regeneration in 35S::THT plants is not driven by the coactivation of multiple hormonal pathways, but rather may rely on a fine-tuned adjustment in auxin levels or distribution.

At the molecular level, *THT* overexpression was associated with a strong up-regulation of *WIND1* in both hypocotyl and cotyledon tissues (Fig. 8E, F). *WIND1* is a central wound-induced transcription factor that promotes cellular dedifferentiation and callus formation, operating at the interface between defence and regeneration (Xu & Yang, 2025). Its elevated expression in *35S::THT* plants indicates that these tissues are transcriptionally activated to regenerate tissues following injury, complementing the enhanced structural reinforcement conferred by HCAA accumulation, callose deposition, and lignification.

Cell wall remodelling is not merely protective but actively instructive during regeneration. In this respect, exogenous cellulose can trigger regenerative responses by integrating defence signalling with hormonal control, positioning the cell wall as a dynamic regulatory platform rather than a passive scaffold (Ruiz-Solaní et al., 2025) Within this framework, the elevated HCAA pool in *35S::THT* plants may contribute to a cell wall environment that balances mechanical reinforcement with controlled plasticity, thereby enabling efficient wound sealing while preserving the capacity for tissue regeneration.

Collectively, these results indicate that THT overexpression enhances regenerative competence after wounding by coupling cell wall remodelling with auxin-associated transcriptional reprogramming, thereby accelerating tissue repair in tomato.

### *THT* overexpression reprograms wound-induced volatile emission and alters volatile-mediated defence signalling

Our results demonstrate that the tomato *THT* gene plays a dual role in plant defence, reinforcing constitutive protection while profoundly altering volatile-mediated communication. Exposure of WT receiver plants to VOCs emitted by wounded WT induced the classical defence marker *PR1* (Fig. 9B), confirming that damage-associated volatiles act as airborne signals capable of activating defence pathways in neighbouring plants. Strikingly, VOCs from non-wounded *35S::THT* plants were sufficient to induce both *PR1* (Fig. 9B) and *TCI21* (Fig. 9A), indicating that THT overexpression promotes constitutive emission of defence-modulating volatiles independently of wounding. Together, these findings indicate that *THT* overexpression alters not only internal metabolic and hormonal states but also the quality and signalling potential of emitted VOCs.

This behaviour is consistent with previous studies showing that plants exposed to volatiles from wounded or stressed conspecifics often upregulate defence-related genes, a phenomenon known as volatile-mediated defence priming (Heil & Karban, 2010; Ángeles-López et al., 2012; Ueda et al., 2012). In tomato, such interplant signalling via VOCs has been associated with enhanced resistance to pathogens and insects, even in the absence of direct damage (López-Ráez et al., 2012). Besides, the volatile bouquet in Solanaceae is highly dynamic and responsive to biotic stress, and alterations in VOC composition can modify both direct and indirect defences (Gargiulo et al., 2025). Our data suggests that *THT* overexpression establishes a constitutive shift in the volatile signature with defence-activating capacity, likely arising from metabolic reprogramming of hydroxycinnamic acid pathways that impacts VOC biosynthesis and release.

To elucidate the basis of this altered signalling capacity, we performed a comprehensive metabolomic analysis. PCA analysis revealed a clear genotype-dependent separation of wounded samples, demonstrating that THT overexpression strongly reprograms the wound-induced volatilome (Fig. 8). In WT tomato plants wounding triggered the emission of classical green leaf volatiles (GLVs) and short-chain C5 compounds such as *Z*-3-hexenal, 1-penten-3-one, Z-3-hexenyl butyrate, and 2,4-hexadienal. GLVs are released under various stress conditions, such as bacterial, fungal, or herbivore attacks (ul Hassan et al., 2015), as well as in response to mechanical damage like wounding (Bate & Rothstein, 1998). These findings confirm that plants respond to wounding through the rapid release of fatty acid-derived volatiles mediated by the *LOX* pathway (Rasulov et al., 2019), which occurs due to damage to the cell membranes. These molecules are rapidly produced after membrane disruption and serve as potent defence signals both within and between plants. In contrast, wounded *35S::THT* plants exhibited a distinct volatile profile enriched in long-chain aldehydes and oxygenated compounds such as E-2-heptenal, E-2-octenal, 2H-pyran-2,6-dione, and 2-pentylfuran, several of which possess documented antimicrobial activity against *Pseudomonas syringae* and other pathogens (López-Gresa et al., 2017; Liarzi et al., 2020; Ma & Johnson, 2021). Interestingly, some of these volatiles, including E-2-heptenal and E-2-octenal, also contribute to the aroma profile of ripe tomato fruit, being associated with characteristic fatty and oily notes (Garbowicz et al., 2018).

In addition to these wound-dependent differences, non-wounded *35S::THT* plants exhibited a differential emission of some monoterpenoids such as β-cymene, D-Limonene, α-terpinene, and also a fatty aldehyde like E-2-nonenal. These compounds were absent or present at much lower levels in the corresponding WT (Fig. 11). Monoterpenoids and fatty aldehydes, play active roles in plant protection against pests, pathogens, and abiotic stress (Gargiulo et al., 2025). In particular, D-limonene and related monoterpenes have been shown to function as both direct defence compounds and airborne signals involved in plant-plant communication (Ben Abdallah et al., 2023; Rosenkranz et al., 2021). Consistent with this, metabolomic profiling of tomato plants infected with *Pseudomonas syringae* revealed that several hydroxylated monoterpenoids are specifically induced as part of the defence response (López-Gresa et al., 2017), and α-terpineol has been shown to exert a direct defensive role against this pathogen (Pérez-Pérez et al., 2024). Moreover, the dynamic nature of VOC blends in tomato has also been demonstrated in response to both pathogenic and beneficial microorganisms: treatments with *Botrytis cinerea* or *Trichoderma virens* altered the emission of monoterpenes, suggesting that VOC output is reprogrammable depending on the biotic context (Nawrocka-Kaczmarek et al., 2023). Therefore, the constitutive emission of β-cymene, α-terpinene, D-limonene, and E-2-nonenal in *35S::THT* plants may reflect a metabolically activated state that enables defensive communication even in the absence of wounding.

Functional assays with individual volatiles support this model. Treatment with *Z-3*-hexenal, a volatile abundant in WT wounded plants, induced *TCI21* expression (Fig. 12D), whereas treatment with a compound characteristic of wounded *35S::THT* plants such as E-2-heptenal failed to elicit any induction of this gene. Consistent with this interpretation, the basal accumulation of HCAAs and the enhanced cell wall fortification observed in these lines likely confer anticipatory protection against wounding, therefore reducing the reliance on inducible volatile-mediated responses. Remarkably, the volatile identified from non-wounded *35S::THT* plants as limonene was able to induce *TCI21*, however without significant differences, in agreement with the interplant assays (Fig. 9), confirming that this genotype constitutively releases compounds capable of priming defence gene expression such as *TCI21* (Fig. 9A) and *PR1* (Fig. 9B). These results establish a causal link between the altered volatilome of *35S::THT* plants and the modulation of defence signalling in neighbouring individuals.

Collectively, our findings identify THT and its hydroxycinnamic acid amide products as a central integrative module in the tomato wound response. By promoting constitutive and wound-induced accumulation of HCAAs, THT reinforces cell wall architecture, stabilises redox homeostasis and enhances regenerative competence without disrupting canonical jasmonate or abscisic acid signalling. At the same time, THT-driven metabolic reprogramming reshapes the volatile blend, shifting defence communication from an inducible, GLV-dominated response towards a constitutively active, HCAA-associated signalling state that modulates defence gene expression in neighbouring plants. Together, these effects reveal how a single phenylpropanoid branch can coordinate local protection, tissue repair and airborne signalling, highlighting HCAAs as multifunctional effectors that couple defence robustness with regenerative capacity during mechanical stress.

## Acknowledgements

We acknowledge the Plant Hormone Quantification Service at the IBMCP (Valencia, Spain) for performing the hormone analyses. We thank Dr. Nuria Coll (Group Leader and CSIC Researcher at Centre for Research in Agricultural Genomics, Barcelona, Spain) for hosting C.G.-H. in her laboratory and for providing expert guidance in microscopy analyses. This work was supported by grants PID2020-116765RB-I00 and PID2023-152361OB-I00 funded by MCIN/AEI/10.13039/501100011033/, also grants by Aid to First Research Projects (PAID-06-25) funded by Vice-Rectorate for Research of thre Universitat Politècnica of Valencia (UPV), and supported by grants PROMETEU/2021/056 and CIPROM/2024/41 from Generalitat Valenciana. C.G.-H is the recipient of a predoctoral fellowship (CEX2021-001172-S-20-8), funded by MCIN/AEI/10.13039/501100011033 and by the European Social Fund Plus (ESF+)

## Author contributions

C.G.-H., C.A., F.V.-S., I.R., J.M.B., M.P.L.-G., and P.L. conceived and designed the study. C.G.-H. and C.A. performed the experiments. C.G.-H., C.A., F.V.-S., I.R., J.M.B., M.P.L.-G., and P.L. analysed and interpreted the data. C.G.-H., M.P.L.-G., F.V.-S., and P.L. wrote and revised the manuscript.

## Data availability

All data is incorporated into the article.

## Supporting Information

**Figure S1.**
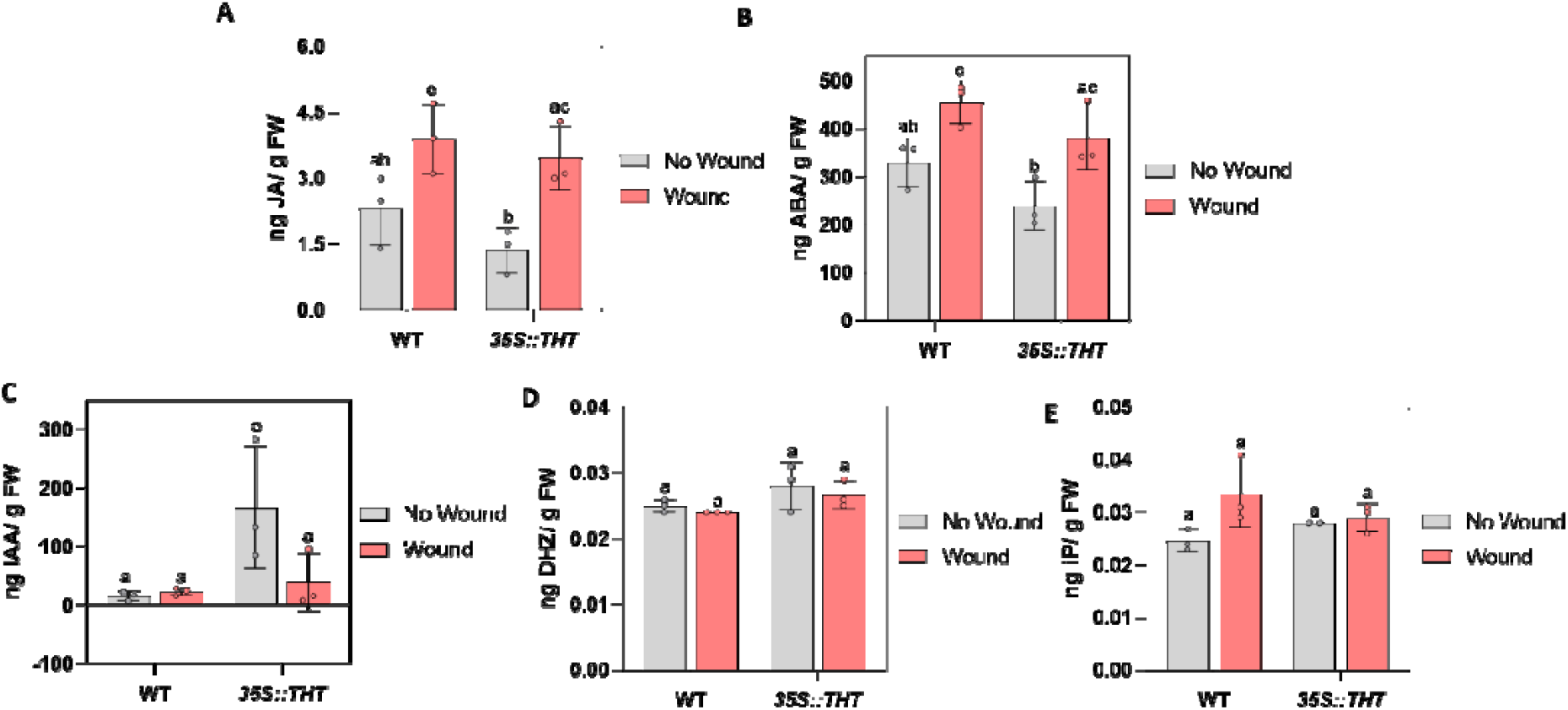
Levels of **A)** jasmonic acid (JA), **B)** abscisic acid (ABA), **C)** indole-3-acetic acid (IAA), **D)** cytokinin dihydrozeatin (DHZ), and **E)** iso-pentenyladenine (iP) in leaves of WT and *35S::THT* plants under non-wounded (No Wound) and wounded (Wound) conditions. Hormone concentrations are expressed as ng·g^−1^ FW and correspond to the mean ± SD of a representative experiment (n = 3). Statistically significant differences (ANOVA, *p* < 0.05) between genotypes and treatment conditions are indicated by different letters.

**Table S1.**
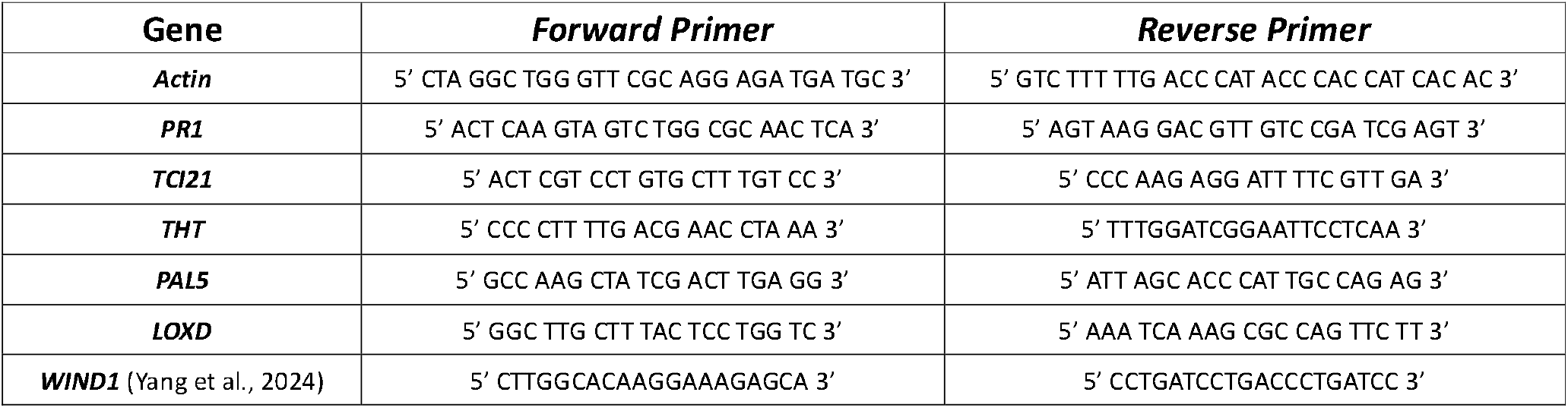
Primer sequences used for RT-qPCR analyses.

